# Multispecies coexistence emerges from pairwise exclusions in communities with competitive hierarchy

**DOI:** 10.1101/2025.04.22.649899

**Authors:** Zachary R. Miller, Dillon Max

**Affiliations:** Department of Earth & Planetary Sciences, Yale University; Department of Ecology & Evolutionary Biology, University of California, Los Angeles

**Keywords:** coexistence, community assembly, emergent coexistence, competitive hierarchy, competition-colonization trade-off, assembly rules

## Abstract

Competitive coexistence is often understood as an additive process where coexisting species pairs, triplets, etc. combine to form larger communities. However, emergent coexistence – where multi-species persistence occurs without pairwise coexistence – can arise through mechanisms including intransitive loops, facilitation, or higher-order interactions. Emergent coexistence has functional consequences, for example constraining community assembly and reducing robustness to extinctions. Here, we demonstrate that emergent coexistence can arise without intransitivity in competitive communities with pairwise interactions. First, we develop a simple trade-off model where interactions are competitive, transitive, and pairwise, yet coexistence is emergent. Second, we show that coexistence is typically emergent in well-known hierarchical trade-off models. Third, we find that emergent coexistence frequently occurs without pronounced intransitivity in random model communities. Our results suggest that competitive coexistence may often be emergent, highlighting a need to better understand the mechanisms and prevalence of this phenomenon in order to reliably predict community assembly, robustness, and biodiversity maintenance.

## Introduction

Coexistence theory plays a central role in explaining the maintenance of biodiversity. This theory is often focused on the coexistence of two or three species, but many natural communities hold tens, hundreds, or even thousands of species. Understanding how coexistence theory scales up to large communities remains a critical question. Modern Coexistence Theory emphasizes the interplay of niche and fitness differences between pairs of species as a unifying framework for coexistence (Chesson, 2000; Barabás *et al*., 2018; Buche *et al*., 2022; Spaak & Schreiber, 2023). However, it has been challenging to extend this framework to larger communities (Saavedra *et al*., 2017; Barabás *et al*., 2018; Chesson, 2018; Song *et al*., 2019; Hofbauer & Schreiber, 2022). More generally, it is rarely clear when traditional characterizations of coexistence – i.e., simple rules based on interversus intraspecific interaction strengths or niche differentiation – apply beyond species pairs to predict coexistence of three or more species (Barabás *et al*., 2016; Levine *et al*., 2017; Miller *et al*., 2022; Ranjan *et al*., 2024).

One natural question is how closely the coexistence of pairs – or other subsets – of species is tied to the coexistence of a larger community they comprise (Engel & Weltzin, 2008; Barabás *et al*., 2016; Saavedra *et al*., 2017). We can imagine two extreme cases: *perfectly additive* coexistence would mean that every subset of a coexisting community also coexists, while *perfectly emergent* coexistence would mean that no subset can persist independently, and coexistence is only possible in the full community. In practice, coexistence may be a more or less emergent property anywhere along this continuum. Pairwise outcomes are of particular interest, since species interactions are usually modeled and measured as pairwise effects. Is coexistence of all species pairs ever necessary or sufficient for coexistence for full community coexistence? While it is often assumed (Adler *et al*., 2018) or explicitly argued (Friedman *et al*., 2017; Ortiz *et al*., 2021) that pairwise interaction outcomes are predictive of full community coexistence – with complex communities assembled in an additive fashion from stable subunits, and pairwise exclusion precluding further coexistence – Chang *et al*. (2023) showed that microbes drawn from stably coexisting communities frequently fail to coexist in isolated pairs. In these systems, coexistence is achieved only in the context of the larger community – coexistence is emergent. Many empirical studies that characterize pairwise niche and fitness differences in putatively coexisting communities find that individual species pairs would not be predicted to coexist (Kraft *et al*., 2015; Buche *et al*., 2022; Yan *et al*., 2022; Pajares-Murgó *et al*., 2024), suggesting that emergent coexistence might be widespread. Moreover, Angulo *et al*. (2021) recently found that “disassembly holes”, where subsets of a coexisting community fail to coexist, are common in both Lotka-Volterra model communities and empirical microbial communities.

Understanding the prevalence and mechanisms of emergent coexistence has significant implications for ecological theory and applications. If coexistence is an emergent property of large communities, the number of paths to assembling the community via sequential species introductions is more limited. As a result, restoring communities through stepwise species introductions (Angulo *et al*., 2021; Serván & Allesina, 2021) or assembling communities from the ground up (e.g. building synthetic microbial communities (Großkopf & Soyer, 2014; Leggieri *et al*., 2021)) may be challenging. Perhaps more importantly, emergent coexistence might imply fragility (Ebenman & Jonsson, 2005), with these communities being prone to secondary extinctions or even collapse when some species are lost (Bascompte & Stouffer, 2009; Brodie *et al*., 2014). From a theoretical standpoint, emergent coexistence suggests a more complex picture of biodiversity maintenance, where pairwise interaction outcomes are insufficient to capture important coexistence mechanisms (Levine *et al*., 2017; Alcántara *et al*., 2017; Aguadé-Gorgorió & Kéfi, 2025).

The concept of emergent coexistence is not new. Coexistence of a set of species without coexistence of all subsets can result from dependencies such as predation, mutualism, and microbial cross-feeding (Ebenman & Jonsson, 2005; Brodie *et al*., 2014; Kehe *et al*., 2021); intransitive competition (Laird & Schamp, 2006; Allesina & Levine, 2011; Soliveres *et al*., 2015); and higher-order interactions (HOIs) (Kelsic *et al*., 2015; Gibbs *et al*., 2024). The potential for these mechanisms to lead to cascading secondary extinctions has been studied in depth (Ebenman & Jonsson, 2005; Eklöf & Ebenman, 2006; Bascompte & Stouffer, 2009; Brodie *et al*., 2014).

Emergent coexistence has received less attention in the context of competitive networks, where it is almost exclusively associated with intransitive competition (Allesina & Levine, 2011; Verdú *et al*., 2023). A classic example of intransitive competition is the rock-paper-scissors motif, a stable three-species model where each species outcompetes and is outcompeted by one other (May & Leonard, 1975; Ranjan *et al*., 2024). Intransitive loops have been engineered in synthetic microbial communities, and can explain why a community of three or more species may coexist together, but not when species are paired in isolation (Kerr *et al*., 2002). When co-occurring species fail to show evidence of pairwise coexistence, intransitive competition is often hypothesized to play a role (Godoy *et al*., 2017; Pajares-Murgó *et al*., 2024). However, the empirical prevalence and importance of competitive intransitivity is unclear. While some studies have indicated a role for competitive intransitivity in facilitating coexistence (Buss & Jackson, 1979; Soliveres *et al*., 2015, 2018; Vandermeer & Perfecto, 2023; Verdú *et al*., 2023), competitive transitivity (hierarchy) also appears common (Savolainen & Vepsäläinen, 1988; Keddy & Shipley, 1989; Grace *et al*., 1993; Shipley & Keddy, 1994; Kinlock, 2019; Koch *et al*., 2023). For example, an analysis of annual plants in a California grassland found that only 15% of species triplets were intransitive, and even then, these relationships were insufficient to stabilize coexistence without pairwise niche differences (Godoy *et al*., 2017). And yet, in the same system, less than 12% of species pairs showed evidence of pairwise coexistence, suggesting that community coexistence may only emerge with more than two species (Kraft *et al*., 2015; Saavedra *et al*., 2017).

The recent experiments by Chang *et al*. (2023) pose an especially sharp challenge to the conventional understanding of emergent coexistence. In these microbial communities, emergent coexistence is widespread even though there are no strong dependencies between species (all but 1 of 62 strains grow in monoculture, suggesting that competitive interactions likely predominate (Ghoul & Mitri, 2016; Kehe *et al*., 2021)) and competitive outcomes form strict hierarchies. These results raise important questions: Is emergent coexistence more prevalent in nature than currently realized? Can mechanisms other than intransitivity cause emergent coexistence in competitive communities? Are HOIs necessary to explain such outcomes?

We consider these questions from a theoretical point of view. We demonstrate that emergent coexistence – defined here as a lack of coexistence among species pairs drawn from a larger community that *does* coexist – can occur in model communities where interactions are strictly competitive, pairwise, and transitive. We first introduce a simple model of a growth-competition trade-off, where emergent coexistence arises from indirect interaction chains. In this model, large communities may contain a vanishing fraction of coexisting pairs. We then show that the conditions leading to emergent coexistence may not be “special” or even uncommon. Indeed, a well-known ecological model, the Hastings-Tilman competition-colonization trade-off model (Hastings, 1980; Tilman, 1994), naturally exhibits both emergent coexistence and competitive hierarchy, with no fine-tuning of the model parameters. We also find that emergent coexistence is common and competitive intransitivity is rarer than expected by chance in model communities with random, unstructured interactions. Together, our results suggest that emergent coexistence may be more prevalent in competitive communities than previously thought and may arise due to unappreciated mechanisms.

## Methods and Results

### Emergent coexistence in a growth-competition trade-off model

To derive a minimal model of emergent coexistence with competitive hierarchy, we consider a version of the generalized Lotka-Volterra (GLV) model constrained by a growth-competition trade-off. By construction, the model does not include positive interactions or HOIs, and any competitive exclusions in species pairs necessarily follow a hierarchy. This model can nevertheless exhibit highly emergent coexistence.

We imagine a pool of *n* species with intrinsic growth rates *r*_*i*_ for all *i* in 1,…, *n*. Species interact through a highly asymmetrical form of interference competition: whenever two individuals encounter one another, the superior competitor kills the inferior (deriving no benefit from the interaction). Competitive ability depends on species identity according to a hierarchy, where species 1 is the best competitor and species *n* is the worst. When two individuals of the same species interact (*i* = *j*), one dies at random. We assume that the disparity in competitive abilities is counterbalanced by an opposing hierarchy of growth rates, so that 0 < *r*_1_ < *r*_2_ < … < *r*_*n*_. This scenario models a trade-off between growth rates and competitive ability due to allocation constraints (Kneitel & Chase, 2004; Garland Jr *et al*., 2022) – for example, competition between bacteria species that can invest resources into either toxin production and defense or multiplication (growth) (Ferenci, 2016).

Assuming the system is well-mixed, its dynamics are described by a GLV model,

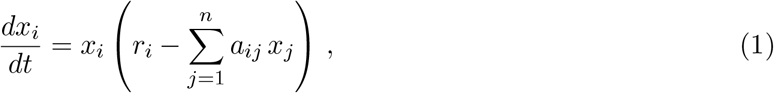

where *x*_i_ is the abundance of species *i* and the matrix of interactions *A* = (*a*_*ij*_) follows the specific form *a*_*ij*_ = 1 if *j*< *i, a*_*ij*_ = 0.5 if *i* = *j*, and zero otherwise.

First, we characterize coexisting communities in this model. Due to the hierarchical structure of the matrix *A*, it is straightforward to prove that any equilibrium of the dynamics is stable (see SI Section A.1). However, in order for all *n* species to have positive abundances at equilibrium (feasibility), the growth rate of each species *i* must exceed a lower limit 𝓁_*i*_ (i.e., *r*_*i*_ > 𝓁_*i*_ for all 1 ≤ *i* ≤ *n*), where 𝓁_*i*_ is defined recursively by

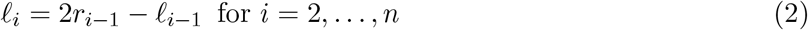

with 𝓁_1_ = 0. We derive these lower limits in SI Section A.2.

Effectively, each species *i* excludes any others with a growth rate in the interval (*r*_*i*_, 𝓁_i+1_). This interval can be visualized as a *niche shadow* cast by species *i* (Fig. 1a; May & Nowak (1994); Kinzig *et al*. (1999)). From Eq. 2, the length of this niche shadow is 𝓁_*i*+1_ − *r*_*i*_ = *r*_*i*_ − 𝓁_*i*_. In other words, proceeding along the growth rate axis starting at *r* = 0, the length of each niche shadow is equal to the distance between the growth rate of the species casting it (*r*_*i*_) and the end of the preceding niche shadow (𝓁_*i*_). In SI Section A.3, we also prove that 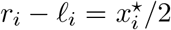, where 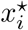 is the equilibrium abundance of species *i*. That is, the length of each species’ niche shadow is proportional to its equilibrium abundance. Intuitively, a species that just barely exceeds its own growth rate threshold (𝓁_*i*_) attains a small abundance at equilibrium and exerts a weak effect on subsequent competitors, while a species that exceeds its threshold by a large amount will attain a large abundance and cast a long niche shadow.

**Figure 1.**
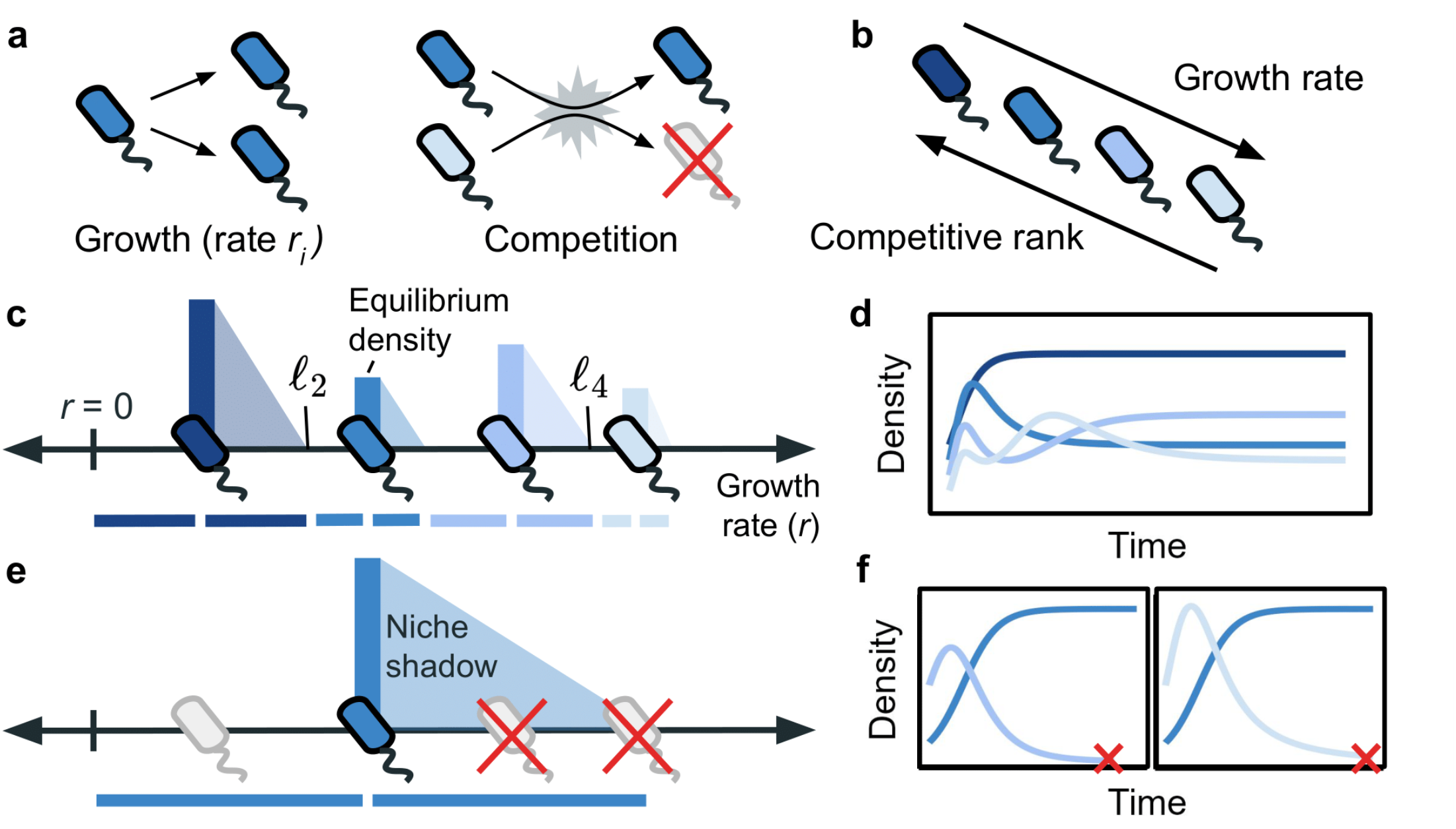
Growth-competition trade-off model. **(a)** Each species *i* grows (reproduces) at a constant rate *r*_*i*_. In bouts of competition, one individual is killed following a competitive hierarchy. **(b)**We assume a growth-competition trade-off, where species that grow faster are worse competitors. **(c)**At equilibrium, each species excludes any others that fall within a “niche shadow” along the trade-off axis (Eq. 2). The length of each shadow is equal to the gap between the species casting it and the previous niche shadow. Species preceded by larger gaps experience less competitive suppression, attain higher equilibrium densities, and cast longer niche shadows (e.g., species 1 and 3 compared to 2 and 4, from left). **(d)** When no species falls in the niche shadow of another, the community dynamics approach a stable equilibrium where all species coexist. **(e)** Removing species 1 allows species 2 (the next best competitor) to increase its niche shadow and exclude species 3 and 4 in pairwise competition **(f)**. Coexistence of species 2, 3, and 4 only emerges in the context of the full community.

We want to understand which and how many species pairs coexist from a stable community of *n* species in this model. The coexistence criteria above suggest a simple way to parameterize all possible coexisting communities. We can specify any positive equilibrium abundances 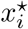, and then define corresponding growth rates

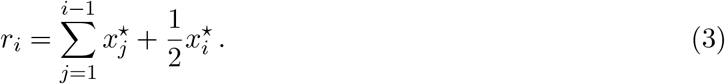

It is clear from Eq. 3 that any feasible equilibrium abundances correspond to a valid sequence of growth rates, with 0 *<* r_1_ < *r*_2_ < … < *r*_*n*_. Notice that the reverse is not true: many choices of growth rates fail to satisfy *r*_*i*_ > 𝓁_*i*_, and therefore do not correspond to a feasible community.

For two species competing in isolation from the rest of the community, the coexistence criteria above become very straightforward: Species *i* and *j > i* coexist if *r*_*j*_ > 2*r*_*i*_. Otherwise, *i* excludes *j*. Three important facts follow: First, any pairwise exclusions respect a hierarchy, with the slower-growing species always excluding the faster-grower if the two cannot coexist. Consequently, competitive outcomes are strictly transitive, with no intransitive loops. Second, pairwise coexistence is most challenging to achieve for adjacent species in the hierarchy; if species *j* coexists with species *j* − 1, then it coexists with all *i<j* − 1, as well. And third, the coexistence of such pairs becomes harder in isolation, compared to the full community context (because *r*_*j*_ must exceed 2*r*_*j*−1_ for the isolated pair to coexist, compared to 2*r*_*j*−1_ − 𝓁_*j*−1_ for coexistence in the full community).

This effect is illustrated visually in Fig. 1e – when species 1 is removed from the community, the abundance of species 2 increases and its niche shadow lengthens, causing species 2 to exclude species 3 and 4 in pairwise competition. Ecologically, we can understand this as a consequence of an interaction chain. For any three species *i* < *j* < *k*, both *j* and *k* experience direct competition from *i*, but the negative effect of *i* on *j* also reduces the competitive pressure exerted on *k* by *j*. The result is an indirect positive effect (often expressed as “the enemy of my enemy is my friend”), which can allow *k* to coexist with *j* in the presence of *i*, even when *j* might exclude *k* in the absence of *i* (Lawlor, 1979; Stone & Roberts, 1991).

Clearly, full community coexistence does not automatically imply coexistence of all species pairs in this model. In fact, coexistence of all pairs in isolation is much more challenging to achieve than coexistence of the *n*-species community they belong to. Coexistence of all adjacent species pairs (which implies coexistence of *all* pairs) demands that *r*_*i*_ > 2*r*_i−1_, or equivalently, 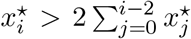.

This requires the sequence of growth rates to increase at least exponentially fast (e.g., Fig. 2, orange) – a much more restrictive condition than the *n*-species coexistence criteria. For example, suppose that we sample coexisting communities by drawing the abundances 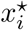 independently from an arbitrary probability distribution. We show in SI Section A.4 that the probability of all pairs coexisting decreases super-exponentially as the size of the community (*n*) grows. For *n* = 10, this probability is already less than 10^−4^.

**Figure 2.**
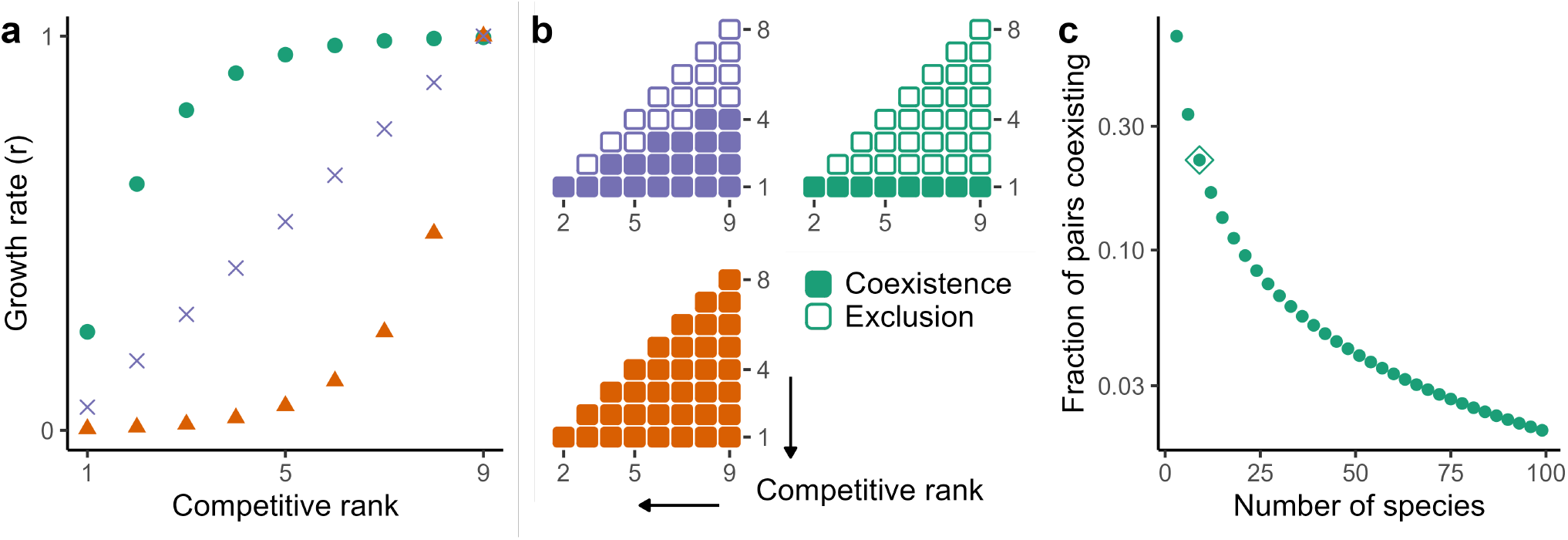
Coexistence fraction depends on trade-off shape. **(a)** Linear (purple), concave (green), and convex (orange) growth rate sequences, all corresponding to coexisting communities of 9 species. **(b)** Pairwise competitive outcomes for all 36 pairs of species in the three communities shown in (a). In the convex community (orange), growth rate increases exponentially, so all pairs coexist. In the concave community (green), *r*_9_ < 2*r*_2_, so all pairs that do not include species 1 result in competitive exclusion. **(c)** If the maximum growth rate *r*_*n*_ is always less than 2*r*_2_ as *n* increases (for example, using 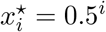 to generate growth rates in Eq. 3), then the coexistence fraction is 2*/n*, which can be arbitrarily small as *n* grows. The community shown in (a) and (b) in green (*n* = 9) is highlighted in (c).

Typically, then, coexistence is to some degree emergent in this model. But what fraction of pairs actually do or do not coexist? Can this coexistence fraction ever be zero, implying a strong form of emergent coexistence?

To answer the second question, we note that the coexistence criterion for species 2 is the same in the full community as in pairwise competition with species 1; in both cases, *r*_2_ must be at least twice *r*_1_. Thus, coexistence of species 1 and 2 in the full community also implies coexistence in isolation. The same is true for species 1 paired with any species *i*, so at least *n* − 1 of the *n*(*n* − 1)*/*2 possible species pairs must coexist.

While this means that the coexistence fraction cannot be zero, it can become arbitrarily small in large communities. Consider that pairwise coexistence of species 2 and *n* requires *r*_*n*_ > 2*r*_2_. If *r*_*n*_ < 2*r*_2_, then species 2 excludes *n* and, moreover, every pair of species 2 ≤ *i<j* will also fail to coexist because *r*_*j*_ *< r*_*n*_ and *r*_*i*_ ≥ *r*_2_. Fig. 2 shows an example. In such communities, the coexistence fraction is 2*/n*, which shrinks to zero as *n* grows (Fig. 2c). In ecological terms, these are communities where species 1 is a very abundant “keystone competitor” (Ennis *et al*., 2023) that equalizes competition between other species pairs; as a result, pairs that do not include species 1 cannot coexist.

To explore the coexistence fraction for parameterizations beyond these extreme cases, we developed a continuum approximation for large communities (SI Section A.5). Briefly, we assume that 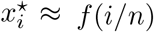 for some continuous function *f*, allowing us to derive an approximate integral formula for the coexistence fraction in arbitrary *n*-species communities.

As an example, this formula can be easily applied in the simple case where species’ abundances follow a power-law: 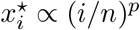 for a constant *p*> −1. For large *n*, the coexistence fraction is approximately 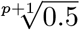. This result reinforces the intuition gained from the extreme cases above: When the trade-off between competitive rank and growth rate is more convex (*p* larger), the coexistence fraction is larger. These are communities where growth rates increase more rapidly with decreasing competitive rank. *p* = 0 corresponds to a community where growth rate varies linearly with competitive rank. In this case, the coexistence fraction is 0.5. For *p*> 0, the sequence of growth rates is convex, and the coexistence fraction is always larger than 0.5. Conversely, if *p <* 0, the trade-off is concave, and the coexistence fraction is smaller. In fact, these results hold in general, for any strictly convex or concave trade-off – not only when the abundances and growth rates follow a power law (see SI Section A.5.2).

Overall, the disruption of interaction chains when species are removed from the community makes pairwise competitive exclusion highly likely in this model. The trade-off can allow the coexistence of any number of species *n*, but coexistence of all pairs further requires a strongly accelerating sequence of growth rates, while any growth rate sequence with *r*_*n*_ < 2*r*_2_ leads to competitive exclusion in nearly all pairs for large *n*.

### Emergent coexistence is typical under trade-offs

Our simple growth-competition trade-off model illustrates that coexistence can be an emergent property in hierarchical competitive communities. However, we might ask whether similar results hold beyond our toy model. Additionally, we have considered communities where growth rates are chosen to allow coexistence. Is emergent coexistence still typical when communities are instead assembled dynamically?

To answer both questions, we turn to the well-known competition-colonization trade-off model developed by Hastings (1980) and Tilman (1994). This is a model of patch dynamics, given by

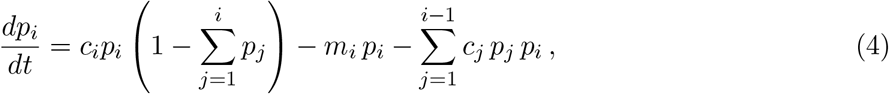

where *p*_*i*_ is the fraction of patches in a landscape that are occupied by species *i* = 1, …, *n*. Species cannot co-occupy patches. The three terms in Eq. 4 correspond, respectively, to colonization of new patches (at rate *c*_*i*_ for species *i*), loss of patches due to disturbance or random local extinction (at rate *m*_*i*_), and loss of patches due to competitive displacement. As in our growth-competition trade-off model, there is a competitive hierarchy – any species *i* can invade and “take over” patches occupied by species *j* > *i*. This is counterbalanced by higher colonization rates or lower mortality rates for worse competitors (i.e., *c*_1_ *< c*_2_ *< … < c*_*n*_ or *m*_1_ > *m*_2_ > … > *m*_*n*_). This model, while still simplistic, reflects trade-offs observed in numerous taxa (Tilman, 1994; Cadotte *et al*., 2006; Rodríguez *et al*., 2007; Yawata *et al*., 2014; Smith *et al*., 2018).

Like our growth-competition above, Eq. 4 can be written as a GLV model (Eq. 1) with growth rate parameters *r*_*i*_ = *c*_*i*_ − *m*_*i*_ and interaction parameters *a*_*ij*_ = *c*_*i*_ + *c*_*j*_ if *j < i, a*_*ij*_ = *c*_*i*_ if *i* = *j*, and zero otherwise. In fact, if all species share a common colonization rate (*c*_*i*_ = *c*), then Eq. 4 becomes exactly equivalent to our growth-competition trade-off model (with the relationship *r*_*i*_ = *c* − *m*_*i*_). This version of Eq. 4 has been studied as a model for infectious disease dynamics with superinfection (Nowak & May, 1994; May & Nowak, 1994), where more competitive strains are also more virulent (*m*_*i*_ *> m*_*j*_ for *i < j*). In this case, our analysis above applies directly, and we predict that strain coexistence due to a competition-virulence trade-off can be highly emergent.

As a model for competition-colonization dynamics in a metacommunity, Eq. 4 has usually been studied under the alternative assumption that local extinction is equal for all species (*m*_*i*_ = *m*), but species differ in colonization ability (*c*_1_ *< c*_2_ *< … < c*_*n*_). In this case, the dynamics are no longer identical to our growth-competition model, but the two models remain qualitatively similar. In particular, pairwise competitive outcomes follow a strict hierarchy, and coexistence can be understood in terms of niche shadows (Kinzig *et al*., 1999), as in our growth-competition model.

Miller *et al*. (2024) showed that with many species (large *n*) the criteria for coexistence in the competition-colonization trade-off model are well-approximated by those found for our growth-competition model (i.e., Eq. 2). This means that full community coexistence can be parameterized similarly using Eq. 3 (see SI Section B for details). For species pairs, on the other hand, the coexistence criteria differ between the two models. Specifically, pairwise coexistence is always harder to achieve in the competition-colonization trade-off model. Thus, one can use our Eq. 3 to construct large, coexisting communities in the competition-colonization model, and these communities will have even fewer pairs of species that coexist in isolation, compared to a growth-competition model with the same growth rates.

Miller *et al*. (2024) used Eq. 2 to study community assembly under the competition-colonization trade-off, starting from a species pool with randomly sampled (and sorted) colonization rates. They found that half of species persist on average in the assembled community, regardless of the size (*n*) or distribution of colonization rates in the species pool. Moreover, the marginal distribution of colonization rates in the assembled community is identical to the “input” distribution in the species pool. Because these conclusions follow from Eq. 2, they also apply to our growth-competition model (if growth rates *r*_*i*_ are sampled randomly).

Together with our analysis in the last section, these facts indicate that coexistence is typically emergent in large, self-assembled communities in both trade-off models. From any random pool of *n* species, a stable community of approximately *n/*2 species assembles, and unless colonization (growth) rate trades off exponentially with competitive rank, a substantial fraction of species pairs will not coexist. If the trade-off is linear (i.e., the distribution of colonization/growth rates in the species pool is uniform), then we would predict that about half of species pairs coexist (or slightly less, in the competition-colonization model). If the trade-off is concave, even fewer pairs coexist in isolation. Notably, the shape of the trade-off strongly impacts the coexistence fraction among pairs, but not the fraction of species that persist in the full community (Miller *et al*., 2024).

Fig. 3 illustrates how large communities assemble in the growth-competition trade-off model (Eq. 1) from a pool of species with random growth rates (a-b, d-e). As predicted, coexistence in these self-assembled communities is highly emergent (c, f). In Fig. 3a-c, growth rates are uniformly distributed; the assembled community has 19 species (from a pool of 40), and only about half of species pairs coexist (44% in this realization). In Fig. 3d-f, the growth rate distribution is slightly concave; the assembled community is similarly large (17 species), and the coexistence fraction among pairs is even lower (16%). Fig. S3 shows very similar results for the competition-colonization trade-off model. In these closely-related trade-off models, assembly from a random pool of species leads to large communities in which coexistence is highly emergent – no fine-tuning of the parameters is needed to produce this property.

**Figure 3.**
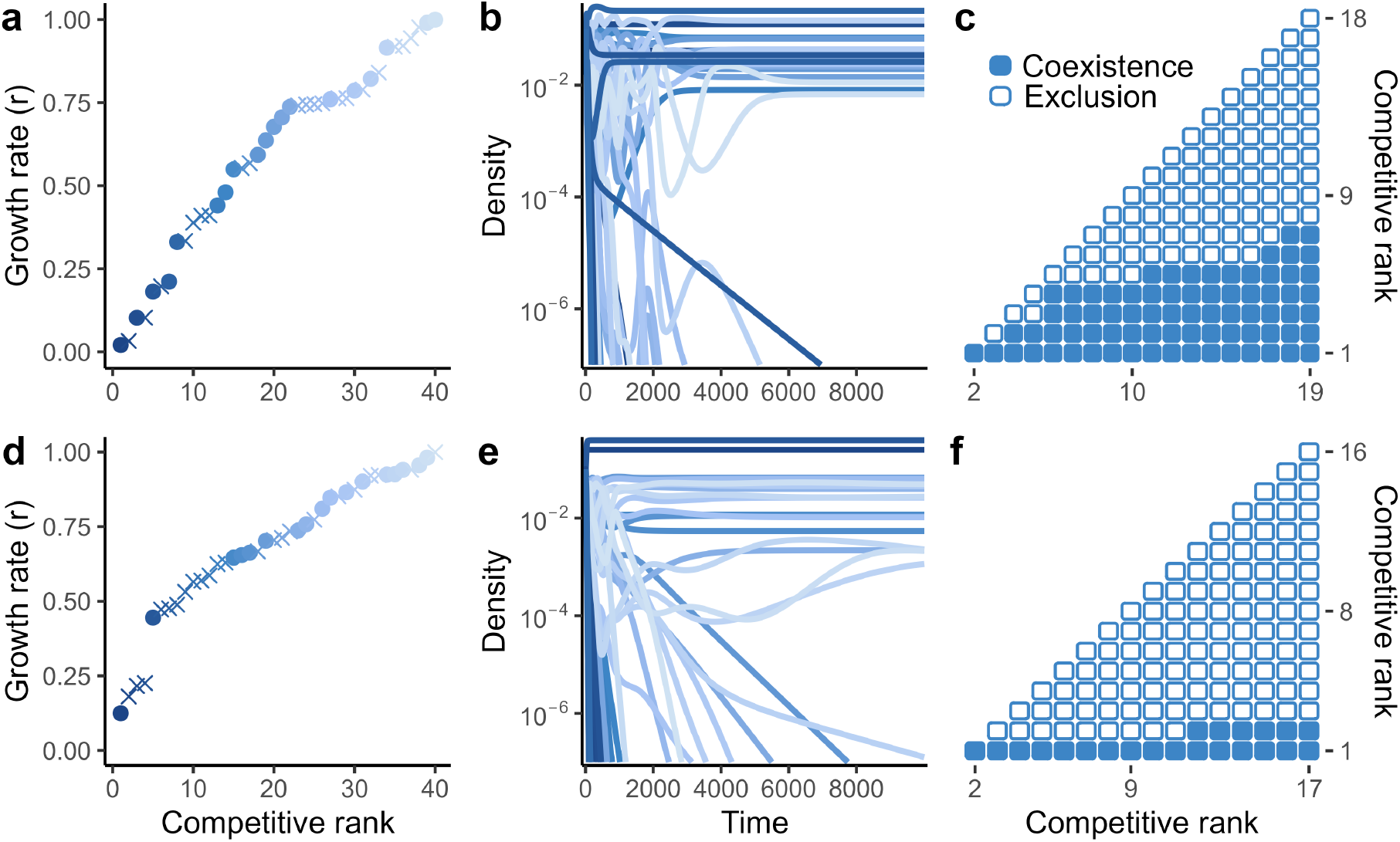
Emergent coexistence in large, self-assembled communities. **(a)** Growth rates are sampled independently from a uniform distribution on [0,1] and sorted so that *r*_1_ *< r*_2_ < … *r*_*n*_. **(b)** For an initial pool of 40 species, we numerically integrated the growth-competition dynamics to obtain a stable community of 19 species (21 species decline to extinction). In (a), persistent species are shown with filled circles, extinct species by crosses. **(c)** A majority of species pairs from the assembled community fail to coexist in isolation. **(d-f)** As in (a-c), except that growth rates are sampled from a triangular distribution (i.e., *P* (*r < ρ*) = *ρ*^2^ on [0,1]), yielding a concave trade-off. Consequently, a smaller fraction of pairs coexist. Fig. S3 shows the corresponding dynamics and pairwise outcomes in the competition-colonization trade-off model (Eq. 4).

### Emergent coexistence and competitive hierarchy without trade-offs

So far, we have examined emergent coexistence in models with strict ecological trade-offs, which allow multispecies coexistence despite a strong pairwise competitive hierarchy. These simple models clearly illustrate how coexistence in the full community can arise through indirect positive interactions, even without pairwise coexistence or intransitive loops. This raises the question of whether highly-structured trade-offs are similarly unnecessary for emergent coexistence. We might expect emergent coexistence more generally whenever pairwise interactions are strongly competitive (favoring pairwise exclusion) but indirect positive effects arise (and promote coexistence) when more than two species interact.

Here, we explore whether emergent coexistence can occur without intransitivity, trade-offs, or other imposed structure in the network of species interactions. To do so, we turn to numerical simulations to study GLV model communities with random competitive interactions. These *in silico* experiments are inspired by the *in vivo* experiments of (Chang *et al*., 2023); mirroring their approach, we constructed coexisting communities with 4-10 species and then evaluated competitive outcomes among all constituent pairs.

To build model communities where *n* species coexisted together but interactions were otherwise as random as possible, we first sampled the competition coefficients *a*_*ij*_ independently from an exponential distribution. Sampling inter- and intraspecific interactions from the same distribution ensured that pairwise competitive effects were strong on average. Next, we sampled (positive) equilibrium abundances 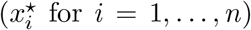 in the same manner. Using the equilibrium relationship 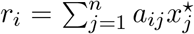 (from Eq. 1) we determined growth rates compatible with these equilibrium abundances. Finally, to ensure the parameters corresponded to a stable community, we evaluated the local stability of the equilibrium by calculating the eigenvalues of the community matrix 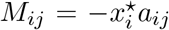. We discarded parameters corresponding to unstable equilibria (those where some eigenvalues had positive real parts), and continued sampling until we had 10^4^ distinct parameterizations for each community size.

Note that the feasibility condition 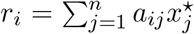 naturally induces a positive correlation between growth rates and interaction strengths (e.g., each species’ mean sensitivity to competition, 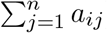. This type of relationship between growth and competition is inevitable in any coexisting GLV community, with the degree of correlation depending on the variance in equilibrium abundances. Our random sampling procedure was designed to minimize this correlation by enforcing a variable species abundance distribution and avoiding any direct trade-offs between growth rates and competitive impacts 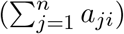 or pairwise outcomes.

For each parameter set, corresponding to a coexisting community of *n* species, we assessed the coexistence of all *n*(*n* − 1)*/*2 pairs in isolation. Although the species belonged to stable *n*-species communities, coexistence of all species pairs was rare, especially in larger communities. For *n>* 5, this occurred in less than 1% of communities, and for *n>* 8, in none. On average, only slightly more than half of pairs coexisted, with no meaningful dependence on community size (mean coexistence fraction of 61% for *n* = 4 and 58% for *n* = 10; see Fig. 4). For every community size, a majority of pairs failed to coexist in at least 25% of communities. Thus, we found that coexistence was often emergent, with coexistence of the full community far from guaranteeing that all, or even most, pairs coexist in isolation.

**Figure 4.**
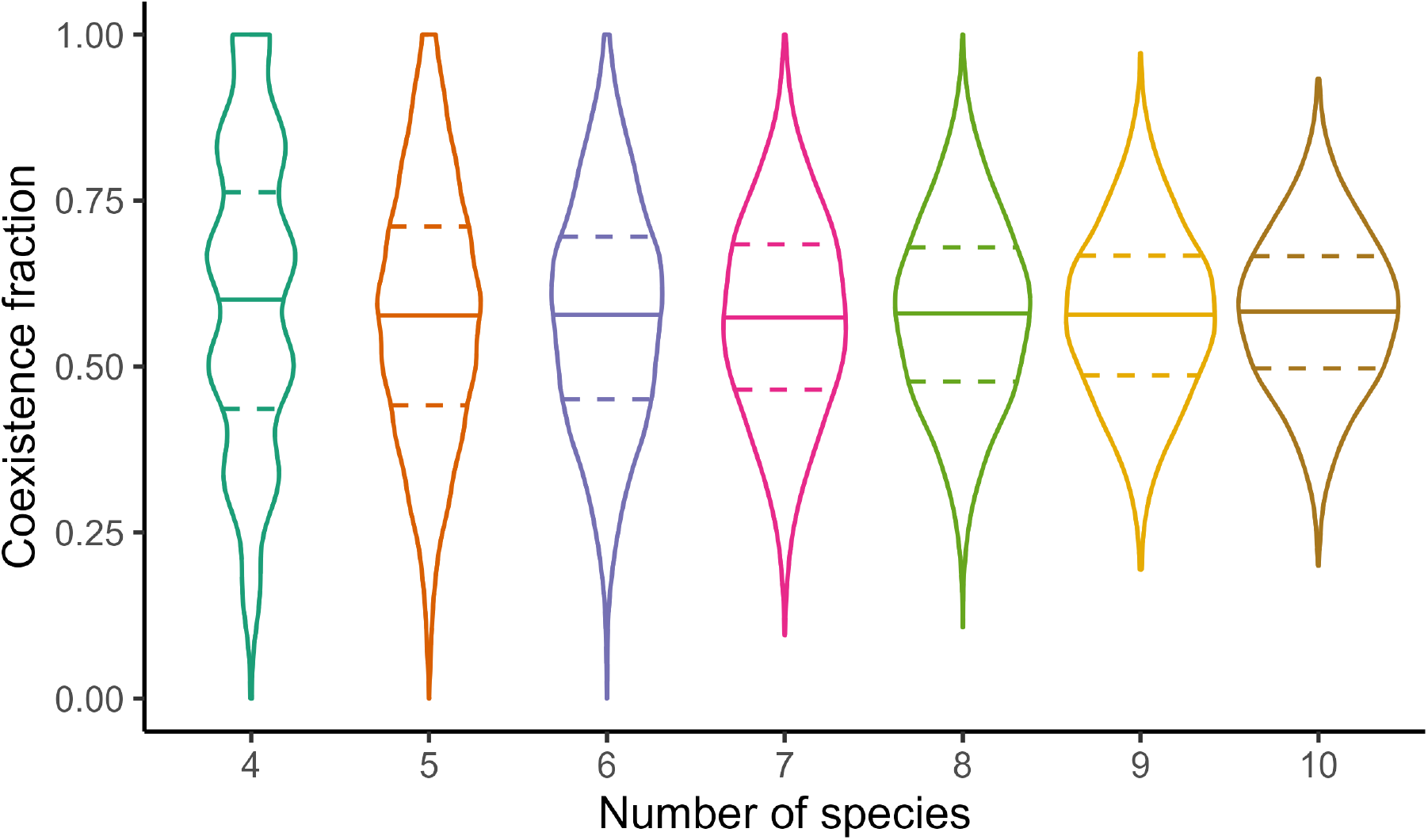
Fraction of species pairs that coexist in random (coexisting) GLV communities. Violins show the density for 10^4^ random communities of 4-10 species (see text for sampling procedure). Medians and quartiles are indicated by solid and dashed lines, respectively. Pairs that do not coexist may exhibit either deterministic competitive exclusion or a priority effect (bistability). The median coexistence fraction does not change substantially as *n* increases, but the distributions become more concentrated as the number of species pairs contributing to the ratio increases, reducing the variance.

Next, we asked whether emergent coexistence in our model communities was associated with competitive intransitivity, as is often assumed when pairwise competitive outcomes fail to predict coexistence.

To assess competitive (in)transitivity, we constructed a network of pairwise outcomes for each community. For every pair of species, we assigned an edge from species *i* to *j* if species *i* excluded *j* in pairwise competition. If the pair coexisted or exhibited a priority effect, we did not assign an edge (priority effects were rare, occurring in less than 5% of pairs for all *n*).

Following Chang *et al*. (2023), we calculated the mean number of intransitive triplets (a set of three species where each excludes exactly one other in pairwise competition) across the competitive networks for all communities. Intransitive triplets represent violations of competitive hierarchy; assessing their prevalence is a common metric of intransitivity (Soliveres *et al*., 2015; Godoy *et al*., 2017; Soliveres *et al*., 2018). We then compared this value to a null distribution, equivalent to randomizing the direction of competitive exclusion (the exact null distribution for the number of intransitive triplets is Binom(*T*, 1*/*4), where *T* is the total number of triplets – across all communities – in which all three pairwise outcomes are competitive exclusions (Chang *et al*., 2023)). For each *n>* 5, the observed mean was smaller than all 10^4^ null model means, indicating under-representation of intransitive triplets in these GLV communities (*n* = 6, 8, and 10 are shown in Fig. 5a; other community sizes shown in Fig. S6).

**Figure 5.**
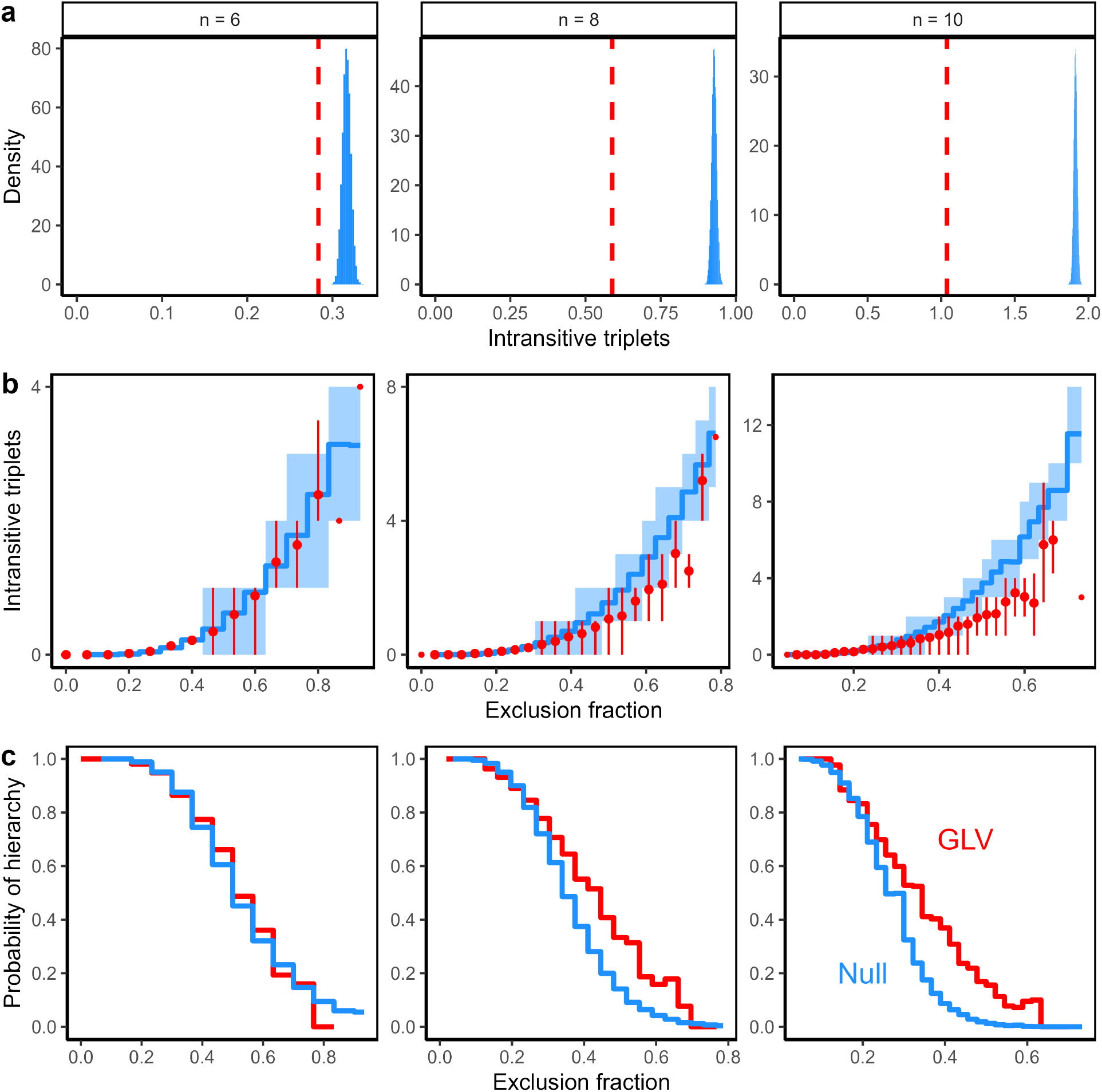
Pairwise competitive outcomes are more transitive than expected by chance in coexisting GLV communities. **(a)** The mean number of intransitive triplets across all 10^4^ replicate GLV communities (red dashed line) compared to a null (binomial) distribution where the direction of competitive exclusion edges is randomly permuted (blue histogram). **(b)** The number of intransitive triplets in GLV simulations (red) vs. a null model (blue), as a function of the exclusion fraction. For each distribution (GLV and null), the mean (red points and blue lines, respectively) and interquartile range (bars and shading) are shown (quartiles not shown for points with < 4 observed communities). **(c)** Fraction of communities with no intransitive loops of any length (GLV simulations in red, null model in blue). As a null model in (b) and (c), we constructed 10^4^ random networks for each observed exclusion fraction where the number of edges was kept the same, but edges were placed and oriented randomly.

While pairwise competitive exclusion is common in this ensemble (Fig. 4) and (triplet) intransitivity is relatively rare (Fig. 5a), one might still wonder whether these properties are positively associated. To investigate this question, we again compared the intransitivity of the competitive networks against null models, but now as a function of the exclusion fraction (i.e., the fraction of pairs exhibiting competitive exclusion). We used a modified null model to account for variation in the exclusion fraction, equivalent to the number of edges in the network. For each unique exclusion fraction observed in our simulated communities, we generated 10^4^ random networks with the same number of edges, but placed and oriented randomly. We then compared the observed versus null model intransitivity using a local metric (number of intransitive triplets) and a global metric (probability of no intransitive loops of any length; i.e., competitive hierarchy). Because the networks have more edges as the exclusion fraction increases, the absolute number of intransitive triplets increases (Fig. 5b) and the probability of hierarchy decreases (Fig. 5c) with the exclusion fraction. However, at any fixed exclusion fraction, we typically find fewer intransitive triplets (Fig. 5b) and a higher probability of hierarchy (Fig. 5c) in the competitive networks, compared to the null model networks. This difference only emerges in larger communities (e.g. *n* = 8 and 10 in Fig. 5) and at higher exclusion fractions, indicating that the communities where coexistence is most emergent are also those where pairwise outcomes are more transitive than expected by chance. Overall, both metrics indicate that intransitivity is not necessary for emergent coexistence in our simulations; if anything, there is a positive association between emergent coexistence (higher exclusion fraction) and more transitive competitive outcomes in larger communities.

## Discussion

It is tempting to account for the coexistence of complex communities in terms of the coexistence of their component species pairs. If coexistence is a simple, additive property, this suggests that large communities are easily assembled from subcommunities and highly robust to species losses. While it is well-understood that coexistence can be more complicated due to ecological dependencies like predation or mutualism (Ebenman & Jonsson, 2005; Brodie *et al*., 2014), intransitivity (Laird & Schamp, 2006; Allesina & Levine, 2011; Soliveres *et al*., 2015), or higher-order interactions (Kelsic *et al*., 2015; Gibbs *et al*., 2024), ecologists often assume that competitive coexistence is essentially additive. Here, however, we have shown that coexistence can be emergent – not reducible to pairwise coexistence – even in competitive communities with hierarchical and strictly pairwise interactions.

We constructed a minimal model with these features and showed that additive coexistence (i.e., co-existence of all pairs) is atypical in this model, while the fraction of coexisting pairs can be arbitrarily small in large coexisting communities. Building on this analysis, we showed that coexistence can also be highly emergent in the well-known competition-colonization trade-off model for metacommunities (Hastings, 1980; Tilman, 1994; Kinzig *et al*., 1999), and we used recent results for that model (Miller *et al*., 2024) to show that coexistence is *typically* emergent in both trade-off models. That is, when large communities self-assemble from a random pool of species, most pairs of species in the final community cannot coexist in isolation. Finally, we used extensive numerical simulations to reveal that emergent coexistence is common in randomly-parameterized communities, conditioned on full community coexistence but without the assumption of trade-offs. Emergent coexistence in these communities is not associated with elevated intransitivity, and perhaps the opposite. Collectively, these results suggest that emergent coexistence may be a general feature of competitive communities (Angulo *et al*., 2021), especially those structured by trade-offs.

Our findings are consistent with a recent theoretical analysis by Aguadé-Gorgorió & Kéfi (2025), who studied emergent coexistence in random GLV models. Aguadé-Gorgorió and Kéfi similarly found that full community coexistence frequently occurs without coexistence of all pairs in a wide parameter regime where interactions are strongly competitive. Our unstructured simulations fall into this regime. Aguadé-Gorgorió and Kéfi argue that emergent coexistence arises when indirect positive effects (via interaction chains) offset strong direct negative interactions, consistent with the mechanism we identify in two trade-off models. In SI Section C, we show that the same factors can explain emergent coexistence in our random GLV simulations. We observe that stronger interspecific competition drives lower pairwise coexistence fractions (Fig. S4) and that our random model communities are enriched in positive indirect effects (compared to a null model where full community coexistence is not enforced; Fig. S5). Like Aguadé-Gorgorió and Kéfi, we find that these indirect effects are linked to intransitive network architectures in the smallest communities (*n ≤* 5) but not in larger communities, where no special structure seems necessary for positive indirect effects to arise. In fact, unlike Aguadé-Gorgorió and Kéfi, we even find that larger communities have less intransitivity than expected by chance – this difference is likely due to the fact that we allow variation in intranspecific competition and growth rates, which promotes competitive asymmetry between species, while Aguadé-Gorgorió and Kéfi only allow variation in interspecific competition strengths.

Understanding whether coexistence is additive or emergent has significant implications for community assembly and robustness. If emergent coexistence is widespread in competitive communities, these systems may be more vulnerable to cascading extinctions than previously thought (Ebenman & Jonsson, 2005; Brodie *et al*., 2014). In communities that are already degraded, restoration from the ground up – through the sequential reintroduction of species – may prove challenging, because fewer assembly paths are viable (Serván & Allesina, 2021). In our trade-off models, for example, assembly can always proceed from best to worst competitor, but starting with any other species can lead to competitive exclusion when a second is added. Similarly, ongoing efforts to engineer synthetic microbial communities for various functional applications (Großkopf & Soyer, 2014; Leggieri *et al*., 2021) are more complicated if emergent coexistence, rather than additive assembly rules (Friedman *et al*., 2017), is typical in complex ecosystems (Angulo *et al*., 2021; Aguadé-Gorgorió & Kéfi, 2025).

Furthermore, our results suggest that patterns of pairwise exclusions can hold relatively little information about full community coexistence and the mechanisms of coexistence. In our minimal model (and thus, in well-known models for competition-colonization (Hastings, 1980; Tilman, 1994) and competition-virulence (May & Nowak, 1994; Nowak & May, 1994) trade-offs), coexistence of a full community is compatible with coexistence of virtually any fraction of species pairs. This underscores the difficulty of making predictions about full community coexistence based on pairwise outcomes, even when full community coexistence is perfectly predictable from *quantitative* knowledge of pairwise interactions (Barabás *et al*., 2016; Alcántara *et al*., 2017; Saavedra *et al*., 2017). Coexistence of all species pairs is known to be insufficient to ensure coexistence of a community (Barabás *et al*., 2016), and our results emphasize that pairwise coexistence is unnecessary, too. Additionally, in both highly-structured trade-off models and unstructured GLV models, we found that emergent coexistence is not indicative of intransitivity, HOIs, or other factors such as environmental fluctuations. Thus, emergent coexistence cannot easily be used to infer that mechanisms beyond pairwise competition are acting in a community.

While our models demonstrate that coexistence can be highly emergent in competitive communities without intransitive loops, we remain far from a general understanding of when competitive coexistence will be additive or emergent. Our results suggest that, in addition to intransitivity, emergent coexistence is also strongly promoted by trade-offs, where a competitive hierarchy at the level of species pairs is counterbalanced by growth, colonization ability, virulence, or other traits, which permit coexistence in the full community context (Hastings, 1980; Tilman, 1994; May & Nowak, 1994; Nowak & May, 1994; Detto *et al*., 2022). However, our numerical simulations imply that intransitivity and trade-offs are not the only mechanisms that can lead to emergent coexistence. These simulation results are consistent with past studies that found extinction cascades (Lundberg *et al*., 2000) and coexistence holes (Angulo *et al*., 2021) – two alternative aspects of emergent coexistence – to be widespread in unstructured model communities. Very recent work by Aguadé-Gorgorió & Kéfi (2025) supports the conclusion that specific interaction structures are unnecessary for emergent coexistence, and makes an important advance by identifying a parameter regime in the random GLV model where emergent coexistence is typical. Interestingly, the analysis of Aguadé-Gorgorió and Kéfi suggests that emergent coexistence will usually be weaker (many, but not all, pairs coexisting) in large (*n>* 10) communities, while we showed that trade-offs can produce highly emergent coexistence even in very large communities.

Additive coexistence, on the other hand, can be expected when interactions are weak (Aguadé-Gorgorió & Kéfi, 2025), structured by phylogeny (Serván *et al*., 2025), or tied to resource competition (Lee *et al*., 2023). However, the last case – resource competition – may be more complex. While Lee *et al*. (2023) found that simple assemble rules assuming additive coexistence are predictive in a mechanistic model of resource competition, Aguadé-Gorgorió & Kéfi (2025) showed that emergent coexistence is also possible in a model based on competition for a single resource plus cross-feeding interactions. These few examples hint that the nature of coexistence, whether additive or emergent, may be tied to the symmetry of competitive interactions. In typical consumer resource models, such as those studied by Lee *et al*. (2023), interactions possess inherent, underlying symmetry, because resource depletion and growth are linked. But this assumption can be violated, leading to asymmetric interactions (Gibbs *et al*., 2022; Liu *et al*., 2025), as in the model developed by Aguadé-Gorgorió & Kéfi (2025). Similarly, intransitive interactions and hierarchical trade-offs both assume strong asymmetries, while models structured by phylogeny are highly symmetric. We hypothesize that asymmetric pairwise interactions are more likely to give rise to strong indirect effects, which are necessary for full community coexistence to be possible when pairwise coexistence is not.

Coupling together pairwise interactions in large ecological communities inevitably gives rise to indirect interaction chains, which can produce emergent outcomes that are challenging to predict from pairwise outcomes alone (Lawlor, 1979; Stone & Roberts, 1991; Wootton, 1994; Saavedra *et al*., 2017; Aguadé-Gorgorió & Kéfi, 2025). While such chains have been studied for decades, it remains necessary to grapple with their consequences for coexistence, robustness, and assembly, especially in ecological communities organized by trade-offs, phylogeny, metabolism, or other large-scale structure. A promising perspective on this problem comes from viewing communities not as a single set of species, but as a nested collection of subcommunities – a community assembly perspective (Song, 2025). Relating full community properties to properties of subcommunities – as we have done here for coexistence in species pairs – has helped shed new light on coexistence (Angulo *et al*., 2021; Hofbauer & Schreiber, 2022; Spaak & Schreiber, 2023), function (Sanchez *et al*., 2023; Arya *et al*., 2023), statistical inference and prediction (Maynard *et al*., 2020), and community change across space and time (Law & Morton, 1996; Law & Leibold, 2005; Serván & Allesina, 2021). In this perspective, emergent coexistence (Chang *et al*., 2023), assembly holes (Angulo *et al*., 2021), and assembly rules (Friedman *et al*., 2017) are fundamental concepts that point to important new dimensions of ecological structure and function.

## Acknowledgments

We thank Maxime Clenet, Stefano Allesina, Chuliang Song, James O’Dwyer, and members of the O’Dwyer lab at the University of Illinois Urbana-Champaign for helpful feedback and insightful discussion.

## Author contributions

ZRM conceived of the research and created the figures. Both authors conducted model development, analysis, and simulations; wrote the text; and contributed to revisions.

## Data accessibility

Code to implement the models and reproduce all simulations and figures is available at https://github.com/zacharyrmiller/emergent_coexistence.

