## Supplementary Information for "Multispecies coexistence emerges from pairwise exclusions in communities with competitive hierarchy"

#### Contents

**A Model analysis** **S2**

A.1 Equilibrium stability in the growth-competition trade-off model . . . . . . . . . . . . . . S2

A.2 Feasibility in the growth-competition trade-off model . . . . . . . . . . . . . . . . . . . . S2

A.3 Derivation of feasible growth rate sequences . . . . . . . . . . . . . . . . . . . . . . . . S3

A.4 Coexistence of all pairs . . . . . . . . . . . . . . . . . . . . . . . . . . . . . . . . . . . S4

A.5 Continuous approximation of the coexistence fraction . . . . . . . . . . . . . . . . . . . . S6

**B Relating the growth-competition and competition-colonization trade-off models** **S9**

**C Emergent coexistence in random GLV communities** **S12**

### A Model analysis

#### A.1 Equilibrium stability in the growth-competition trade-off model

Our growth-competition trade-off model can be written as a generalized Lotka-Volterra (GLV) model, which takes the form:

$$\frac{dx_i}{dt} = x_i \left( r_i - \sum_{j=1}^{n} a_{ij}\, x_j \right) \tag{S1}$$

where $x_i$ is the abundance of species $i$ and the matrix of pairwise interactions $A = (a_{ij})$ follows the specific form $a_{ij} = 1$ if $j < i$, $a_{ij} = 1/2$ if $i = j$, and zero otherwise. Let us assume that $x^\star$ is a feasible equilibrium of Eq. S1 (i.e. $x_i^\star > 0$ and $\frac{dx_i}{dt} = 0$ for each $i = 1 \dots n$). Hereafter, we always take $x$ to be at equilibrium and drop the "star" notation. At equilibrium, Eq. S1 implies

$$r_i = \sum_{j=1}^{n} a_{ij}\, x_j\,. \tag{S2}$$

We want to show that this equilibrium is stable, regardless of the particular population sizes. The Jacobian matrix from Eq. S1 is

$$\begin{aligned} J &= \begin{bmatrix} r_1 - x_1 & 0 & 0 & \dots & 0 \\ -x_2 & r_2 - x_1 - x_2 & 0 & \dots & 0 \\ -x_3 & -x_3 & r_3 - \sum_{j=1}^{3} x_j & \dots & 0 \\ \vdots & \vdots & \vdots & \ddots & \vdots \\ -x_n & -x_n & -x_n & \dots & r_n - \sum_{j=1}^{n} x_j \end{bmatrix} \\ &= \begin{bmatrix} -\frac{1}{2}x_1 & 0 & 0 & \dots & 0 \\ -x_2 & -\frac{1}{2}x_2 & 0 & \dots & 0 \\ -x_3 & -x_3 & -\frac{1}{2}x_3 & \dots & 0 \\ \vdots & \vdots & \vdots & \ddots & \vdots \\ -x_n & -x_n & -x_n & \dots & -\frac{1}{2}x_n \end{bmatrix} \end{aligned} \tag{S3}$$

where each $x_i$ is taken to be the equilibrium value (i.e., $J$ is the community matrix). Because the equilibrium is feasible, the diagonal entries are all negative. And because the matrix is triangular, the diagonal entries are its eigenvalues. Thus, the equilibrium is asymptotically stable.

More generally, if we consider the restriction of Eq. S1 to a subset of species with a feasible equilibrium, we find that the equilibrium is stable. Thus, coexistence in this model is determined solely by feasibility (and invasibility).

#### A.2 Feasibility in the growth-competition trade-off model

We aim to characterize communities with a feasible equilibrium, as a function of the growth rates $r_i$ for $i$ in $1 \dots n$. We will show that there is a lower limit, $\ell_i$, which each growth rate $r_i$ must exceed in order for all species to coexist. Moreover, these lower limits can be defined recursively, leading to the understanding of niche shadows that we develop in the main text.

We start with the following proposition:

**Proposition A.2:** For each species, $i > 1$, the equilibrium population size $x_i$ is non-zero if and only if its growth rate $r_i$ satisfies

$$r_i > 2(r_{i-1} - r_{i-2} + r_{i-3} - \cdots \pm r_1)\,. \tag{S4}$$

*Proof of Proposition A.2*: In order to prove the claim in Proposition A.2, we establish that:

$$\frac{1}{2}x_i = r_i - 2(r_{i-1} - r_{i-2} + r_{i-3} - \cdots \pm r_1) \tag{S5}$$

at equilibrium, from which the claim follows directly.

We derive Eq. S5 inductively. It is clear from Eq. S2 that, at equilibrium, $x_1/2 = r_1$ and $x_2/2 = r_2 - x_1 = r_2 - 2r_1$. We now suppose that Eq. S5 holds for all $j$ up to some $i \geq 2$. From Eq. S2, we have

$$\frac{1}{2}x_{i+1} = r_{i+1} - \sum_{j=1}^{i} x_j\,. \tag{S6}$$

Applying Eq. S5 for each $x_j$ in the sum, we find, after canceling terms, that

$$\frac{1}{2}x_{i+1} = r_{i+1} - 2(r_i - r_{i-1} + \cdots \pm r_1)\,. \tag{S7}$$

Thus, by induction, Eq. S5 holds for every $i \geq 1$ and the proposition follows.

*Recursive lower limits:* From Eq. S4, feasibility requires that $r_i > \ell_i$ for all $i = 1 \dots n$ with $\ell_i$ given explicitly by

$$\ell_i = 2(r_{i-1} - r_{i-2} + r_{i-3} - \cdots \pm r_1)\,. \tag{S8}$$

Alternatively, $\ell_i$ can be defined recursively by

$$\ell_i = 2r_{i-1} - \ell_{i-1} \tag{S9}$$

with $\ell_1 = 0$. Eq. S9 is obtained by adding $\ell_{i-1}$ and $\ell_i$ using Eq. S8:

$$\ell_i + \ell_{i-1} = 2(r_{i-1} - r_{i-2} + r_{i-3} - \cdots \pm r_1) + 2(r_{i-2} - r_{i-3} + - \cdots \pm r_1) = 2r_{i-1}\,, \tag{S10}$$

and thus $\ell_i = 2r_{i-1} - \ell_{i-1}$. No species with $r \in (r_i, \ell_i)$ can invade or persist in the community; we refer to the intervals $(r_i, \ell_i)$ as the niche shadows associated to each species $i$.

#### A.3 Derivation of feasible growth rate sequences

In order to study the fraction of species pairs that coexist in large communities, we want a simple way to parameterize coexisting communities of size $n$. Here, we show that we can specify any coexisting community simply by specifying a set of positive equilibrium abundances.

Combining Eq. S5 and Eq. S8, we see that

$$\frac{1}{2}x_i = r_i - \ell_i\,. \tag{S11}$$

This gives us a simple relationship between growth rates $r_i$, lower limits $\ell_i$, and equilibrium abundances, $x_i$. Given a sequence of $x_i$, we can easily reconstruct the corresponding $r_i$ using Eq. S9. Alternatively, we can use the equilibrium condition (Eq. S2) directly:

$$r_i = \sum_{j=1}^{i-1} x_j + \frac{1}{2} x_i \,. \tag{S12}$$

Using this definition, $r_1 = x_1/2 > 0$, and $r_i - r_{i-1} = (x_i + x_{i-1})/2 > 0$, meaning that the sequence $r_1, r_2, \ldots r_n$ is positive and increasing, as required by the assumptions of the model.

#### A.4 Coexistence of all pairs

Given a feasible community, all pairs of species coexist in isolation if and only if $r_j > 2r_i$ for all $j > i$. It is sufficient to consider whether all adjacent pairs of species coexist, which requires $r_i > 2r_{i-1}$ for all $i > 1$. Parameterizing the community in terms of abundances, we have the equivalent requirement

$$\sum_{j=1}^{i-1} x_j + \frac{1}{2} x_i > 2 \sum_{j=1}^{i-2} x_j + x_{i-1} \tag{S13}$$

or, simply,

$$x_i > 2 \sum_{j=1}^{i-2} x_j \text{ for all } i > 2. \tag{S14}$$

This is a very stringent condition. If we imagine sampling the abundances $x_i$ independently from some probability distribution, the probability of satisfying Eq. S14 depends on the specific distribution. However, a necessary (but far from sufficient) condition for coexistence of all pairs is that $x_1 < x_3 < x_5 < \ldots$ and $x_2 < x_4 < x_6 < \ldots$. This is a straightforward consequence of Eq. S14. Sampling abundances independently, the probability of obtaining these specific orderings is

$$\frac{1}{(\lceil n/2 \rceil)!(\lfloor n/2 \rfloor)!} \tag{S15}$$

(where $\lceil x \rceil$ and $\lfloor x \rfloor$ are the ceiling and floor functions, respectively). Eq. S15 provides an upper bound on the probability of all pairs coexisting. Using Stirling's approximation for the factorial, this upper bound is well-approximated by $(\pi n)^{-1}(n/2e)^{-n}$ which decreases super-exponentially. Thus, we expect coexistence of all pairs to be extremely rare for even moderate $n$, even when sampling from the space of feasible full communities.

Note that this upper bound derives from the strict constraints on the ordering of the $x_i$. Additionally, the $x_i$ must generally increase exponentially in order to satisfy Eq. S14. In comparison, to sample a community with the maximal exclusion fraction (i.e., where no pairs coexist except those including species 1), the abundances must similarly vary exponentially, but there is only a weak constraint on their order ($x_1$ must be the largest abundance). For this reason, we expect maximal exclusion to be much more common than maximal (i.e. complete) pairwise coexistence when the abundances are sampled randomly. In Fig. S1, we test this prediction by sampling $10^7$ sets of abundances independently and identically from an exponential distribution; for each random community, we check whether all pairs coexist (Eq. S14 is satisfied), whether the maximal exclusion fraction is realized ($r_n < 2r_2$), or neither. The probability of each of these outcomes is shown for $n$ between 4 and 15. Both extreme outcomes become rarer as $n$ increases, but coexistence of all pairs is much less likely at every $n$ (and this probability declines much faster).

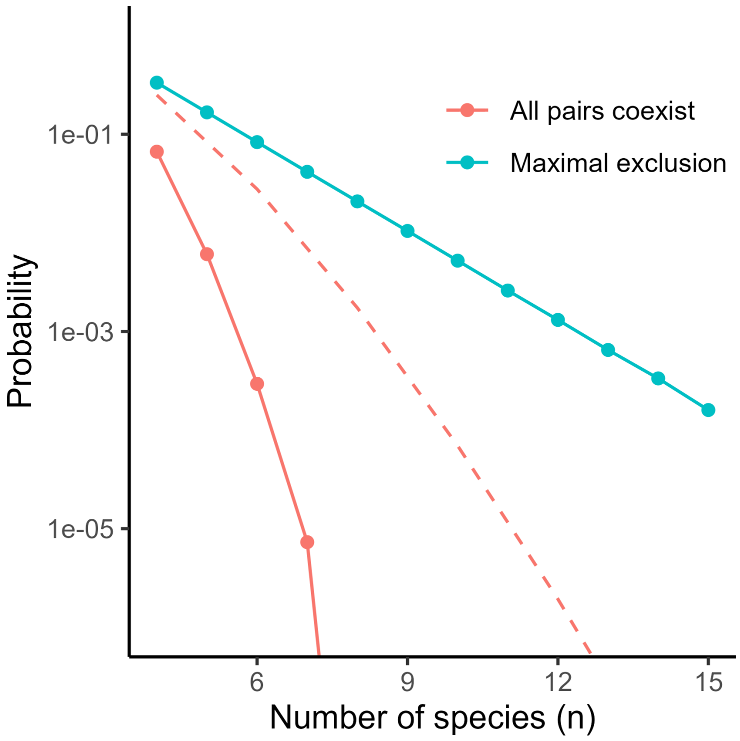


Figure S1: **Probability of complete pairwise coexistence versus maximal pairwise exclusion.** For $n$ between 4 and 15, we sampled $10^7$ parameterizations of the growth-competition model, where coexisting communities of $n$ species were constructed by sampling equilibrium abundances independently from an exponential distribution (i.e., $x_i \overset{iid}{\sim} \text{Exp}(1)$ for all $i = 1 \ldots n$). For each set of parameters, we checked whether all pairs of species could coexist in isolation (requiring $x_i > 2\sum_{j=1}^{i-2} x_j$ for all $i = 1 \ldots n$) or whether all pairs except those involving species 1 resulted in competitive exclusion (requiring that $x_1 > 2\sum_{j=1}^{n-1} x_j + x_n$). The latter scenario is the maximal amount of pairwise exclusion, because pairs involving species 1 necessarily coexist. The probability of these two "extrema" outcomes (across the $10^7$ random communities) are plotted in red and blue, respectively. Maximal exclusion is much more likely than complete pairwise coexistence, and the probability of complete pairwise exclusion decreases super-exponentially with $n$ (compared to an exponential decrease for the probability of maximal pairwise exclusion). An upper bound for the probability of complete pairwise coexistence (Eq. S15) is shown as a dashed line. This loose upper bound is valid for any choice of the equilibrium abundance distribution (provided that abundances are sampled independently).

##### A.5 Continuous approximation of the coexistence fraction

To better understand how the distribution of abundances – or, equivalently, the shape of the growth rate trade-off – affects the fraction of pairs that coexist in isolation, we develop a continuous approximation for this quantity. In an $n$-species community, the growth rates follow:

$$r_i^{(n)} = 2\sum_{j=1}^{i-1} x_j^{(n)} + x_i^{(n)}\,. \tag{S16}$$

We suppose the abundances $x_i^{(n)}$ can be interpolated by a continuous function $f(i,n)$, such that we have

$$r_i^{(n)} = 2\sum_{j=1}^{i-1} f(j,n) + f(i,n) \tag{S17}$$

Now we imagine a sequence of larger and larger communities in which $r_i$ is well-approximated by $r(i/N)$ for some fixed function $r(y)$ as $N$ grows. In order to satisfy these assumptions, we take $f(i,N)$ to be of the form

$$f(i,N) = \frac{1}{N} g\left(\frac{i}{N}\right) \tag{S18}$$

for some fixed function $g(y)$, with $y \in (0,1)$. Now we have

$$r_i = r\left(\frac{i}{N}\right) = 2\sum_{j=1}^{i-1} \frac{1}{N} g\left(\frac{j}{N}\right) + \frac{1}{N} g\left(\frac{i}{N}\right)\,. \tag{S19}$$

Interpreting the sum as a Riemann sum, we can obtain the integral approximation

$$r\left(\frac{i}{N}\right) = 2\int_{\frac{1}{N}}^{\frac{i-1}{N}} g(s)\,ds + \frac{1}{N} g\left(\frac{i}{N}\right) \tag{S20}$$

for large $N$. Letting $y = \frac{i}{N}$ and neglecting $1/N$ terms, we are left with

$$r(y) = 2\int_0^y g(s)\,ds \tag{S21}$$

as a continuous approximation for the growth rates.

Now we consider the pairwise coexistence criterion in this setting. Two species $i > j$ with $i/n \approx y$ and $j/n \approx z$ will coexist if $r(y) > 2r(z)$, or equivalently, if

$$\int_0^y g(s)\,ds > 2\int_0^z g(s)\,ds \tag{S22}$$

The boundary between coexistence and non-coexistence is the curve where $r(y) = 2r(z)$. We denote this curve $z = h(y)$, defined implicitly by

$$\int_0^y g(s)\,ds = 2\int_0^{h(y)} g(s)\,ds\,. \tag{S23}$$

From $h(y)$, we can calculate the (approximate) coexistence fraction, $C$:

$$C = 2\int_0^1 h(y)\,dy \tag{S24}$$

The factor of 2 here comes from the fact that we are integrating over the half-domain $y > z$ (symmetric to the region $y < z$).

We note that the $n-1$ pairs including species 1 must always coexist (as discussed in the main text). Thus, a somewhat better approximation for the coexistence fraction (which captures the dependence on $n$) is given by the "adjusted" approximation

$$\tilde{C} = \frac{2}{n} + \left(1 - \frac{2}{n}\right) C\,. \tag{S25}$$

##### A.5.1 Example: coexistence fraction with equal abundances

To illustrate the use of the formula above, we consider the simple case where all species have equal abundances at equilibrium. This means that $g(y) = c$ is a constant function, and the sequence of growth rates $r_i$ increases linearly. In this setting, we can solve for $h(y)$ explicitly:

$$\int_0^y c\,ds = 2\int_0^z c\,ds \;\Rightarrow\; cy = 2cz \tag{S26}$$

which gives us the equation $z = h(y) = \frac{1}{2}y$. Then we have

$$C = 2\int_0^1 \frac{1}{2}y\,dy = \frac{1}{2} \tag{S27}$$

and

$$\tilde{C} = \frac{2}{n} + \left(1 - \frac{2}{n}\right)\frac{1}{2} = \frac{1}{2} + \frac{1}{n} \tag{S28}$$

##### A.5.2 Convex and concave trade-offs

In the simple example above, it is possible to solve Eq. S23 and obtain a closed-form expression for $h$ and subsequently for the coexistence fraction. In general, this may not be possible. However, we can still learn something about the coexistence fraction from this analysis when the sequence of species abundances $x_i$ is strictly increasing or decreasing (as a function of decreasing competitive rank). Equivalently, this means that the sequence of growth rates is strictly convex (accelerating) or concave (decelerating), respectively.

If $g$ is increasing (meaning $r$ is convex), then for every $y$ we must have $h(y) > \frac{1}{2}y$, because

$$\int_{y/2}^{y} g(s)\,ds > \int_0^{y/2} g(s)\,ds\,. \tag{S29}$$

By comparison to the case where $g$ is constant, we conclude that $C > \frac{1}{2}$. If $g$ is decreasing ($r$ concave), then these inequalities are all reversed.

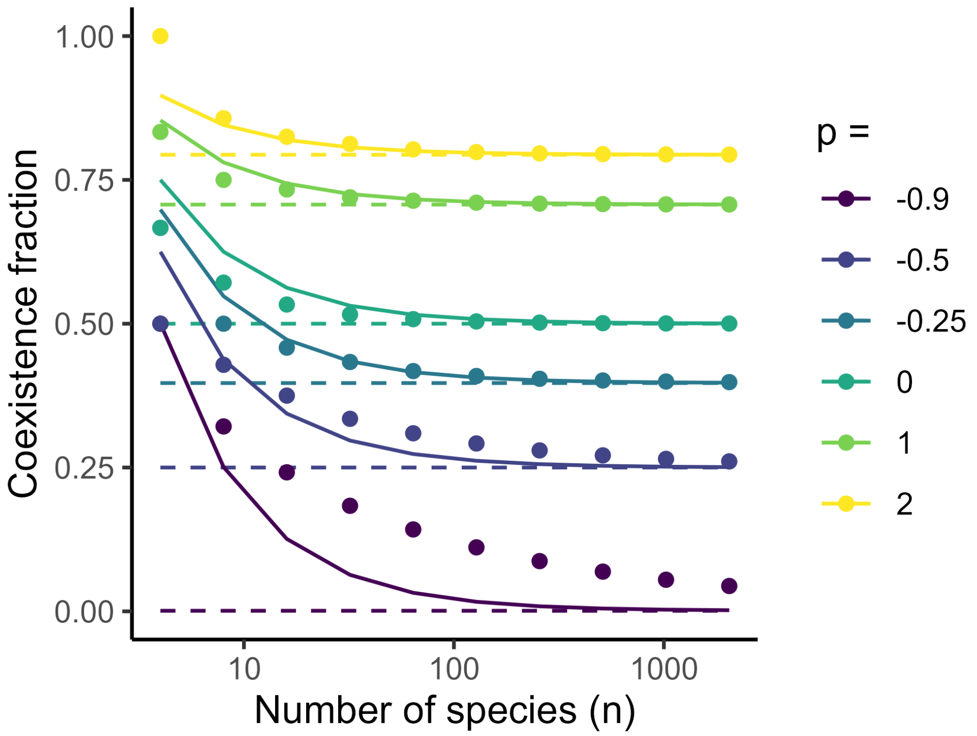


Figure S2: **Validation of the coexistence fraction approximation for power-law communities**. We plot the exact pairwise coexistence fraction (dots) and continuous approximations (based on Eqs. S24 and S25; dashed and solid lines, respectively) for communities generated according to $x_i = (i/n)^p$ for different values of $p$ (colors). Our adjusted approximation (Eq. S25) provides a reasonable match to the dependence on $n$, while our asymptotic (Eq. S31, based on Eq. S24) is accurate for sufficiently large $n$.

##### A.5.3 Example: coexistence fraction with a power-law abundance distribution

It is possible to compute the coexistence fraction whenever the sequence of species abundances follows a power law: $g(s) = cs^p$ for some $c > 0$ and $p > -1$. We solve

$$c \int_0^y s^p ds = 2c \int_0^z s^p ds \Rightarrow \frac{c}{p+1} y^{p+1} = \frac{2c}{p+1} z^{p+1} \tag{S30}$$

yielding $h(y) = \sqrt[p+1]{0.5}y$. From Eq. S24, we have

$$C = 2 \int_0^1 \sqrt[p+1]{0.5}\, y \, dy = \sqrt[p+1]{0.5} \, . \tag{S31}$$

This family of trade-offs illustrates the general conclusions above: For $p > 0$, $g$ is increasing ($r$ is convex), and the predicted coexistence fraction is greater than $\frac{1}{2}$. For $p < 0$, $g$ is decreasing ($r$ is concave), and the coexistence fraction is less than $\frac{1}{2}$. With $p = 0$, we recover the linear case. Notably $C < 1$ for any finite choice of $p$, indicating that coexistence of all pairs is incompatible with any power-law distribution of abundances (for large $n$).

### B Relating the growth-competition and competition-colonization trade-off models

Our growth-competition trade-off model is closely related to the well-known Hastings-Tilman competition-colonization trade-off model (Hastings, 1980; Tilman, 1994):

$$\frac{dp_i}{dt} = c_i p_i \left(1 - \sum_{j=1}^{i} p_j\right) - m_i\, p_i - \sum_{j=1}^{i-1} c_j\, p_j\, p_i\,, \tag{S32}$$

where $p_i$ is the fraction of patches in a landscape that are occupied by species $i = 1 \ldots n$. The parameters $c_i$ are colonization rates and $m_i$ are local extinction rates. An interpretation of the model is given in the main text.

These dynamics can be written as a GLV model with growth rate parameters $r_i = c_i - m_i$ and interaction parameters $a_{ij} = c_i + c_j$ if $j < i$, $a_{ij} = c_i$ if $i = j$, and zero otherwise. As noted in the main text, if $c_i = c$ for all species and $m_1 > m_2 > ... > m_n$, then the competition-colonization trade-off model becomes formally equivalent to our growth-competition trade-off model (up to a re-scaling of all parameters by $c$, which has no effect on species coexistence).

More typically, the competition-colonization trade-off is studied under the assumption that $m_i = m$ for all species, with $c_1 < c_2 < \ldots c_n$. For this formulation of the model, coexistence of all species requires that $c_i > \ell'_i$ for all $i$, with $\ell'_i$ defined recursively by

$$\ell'_i = \frac{c_{i-1}^2}{\ell'_{i-1}} \tag{S33}$$

with $\ell'_1 = m$. Assuming that the $c_i$ are sampled randomly (and independently) from a fixed distribution, Miller *et al.* (2024) noted that $c_{i-1}^2$ can be expanded as $(c_{i-1} - \ell'_{i-1} + \ell'_{i-1})^2 = (c_{i-1} - \ell'_{i-1})^2 + 2\ell'_i(c_{i-1} - \ell'_{i-1}) + \ell'^2_{i-1}$. With many species (large $n$), the quantities $c_{i-1}$ and $\ell'_{i-1}$ are both at least $m$, while $c_{i-1} - \ell'_{i-1}$ shrinks to zero as more species are packed into the species pool. Thus, Eq. S33 can be well-approximated by neglecting the small quadratic term:

$$\ell'_i = \frac{(c_{i-1} - \ell'_{i-1})^2 + 2\ell'_i(c_{i-1} - \ell'_{i-1}) + \ell'^2_{i-1}}{\ell'_{i-1}} \approx \ell'_{i-1} + 2(c_{i-1} - \ell'_{i-1}) = 2c_{i-1} - \ell'_{i-1}\,. \tag{S34}$$

Finally, we can use the change of variables $r_i = c_i - m$ and $\ell_i = \ell'_i - m$ to see that coexistence (approximately) requires $r_i > \ell_i$ with

$$\ell_i = 2r_{i-1} - \ell_{i-1} \tag{S35}$$

and $\ell_1 = 0$, exactly as for our growth-competition trade-off model. Thus, in the limit of large $n$, we expect community assembly to proceed almost identically in these two models; given a pool of species characterized by growth rates $r_i$, the set of species that persist at equilibrium will be almost the same between the two models.

It is important to note that this convergence relies on large $n$; therefore, it does not apply to isolated species pairs drawn from the full community. For two species $j > i$ in the competition-colonization trade-off model, coexistence requires that $c_j > \frac{c_i^2}{m}$. Meanwhile, in the growth-competition trade-off model, we have the condition $r_j > 2r_i$. Re-writing the latter inequality in terms of colonization and local extinction rates, we have $c_j - m > 2(c_i - m)$ or $c_j > 2c_i - m$. Now we can see that

$$\frac{c_i^2}{m} - (2c_i - m) = \left(\frac{c_i}{\sqrt{m}} - \sqrt{m}\right)^2 > 0\,, \tag{S36}$$

implying that the condition for pairwise coexistence in the competition-colonization trade-off model ($c_j > \frac{c_i^2}{m}$) is always more stringent than the corresponding condition in the growth-competition trade-off model ($c_j > 2c_i - m$).

Overall, these results tell us that community assembly from a large random pool will be nearly identical for both models. From Miller *et al.* (2024), this means that approximately half of species will persist at equilibrium, and the marginal distribution of growth rates of these species will be unchanged from the distribution in the pool. From our analysis of the growth-competition trade-off model, we can approximate the coexistence fraction among species pairs given the distribution of growth rates. We have seen that, in the growth-competition model, this fraction will be less than one unless the sequence of growth rates increases exponentially fast. And we have just shown that the coexistence fraction in the competition-colonization trade-off model – assuming the same set of species persist at equilibrium in the full community – will generally be even lower.

These results are illustrated for the growth-competition trade-off model in Fig. 3 (main text). In Fig. S3, we show the corresponding plots for the competition-colonization trade-off model (i.e., the outcome of assembly and pairwise competitions using the same pool of growth rates $r_i$).

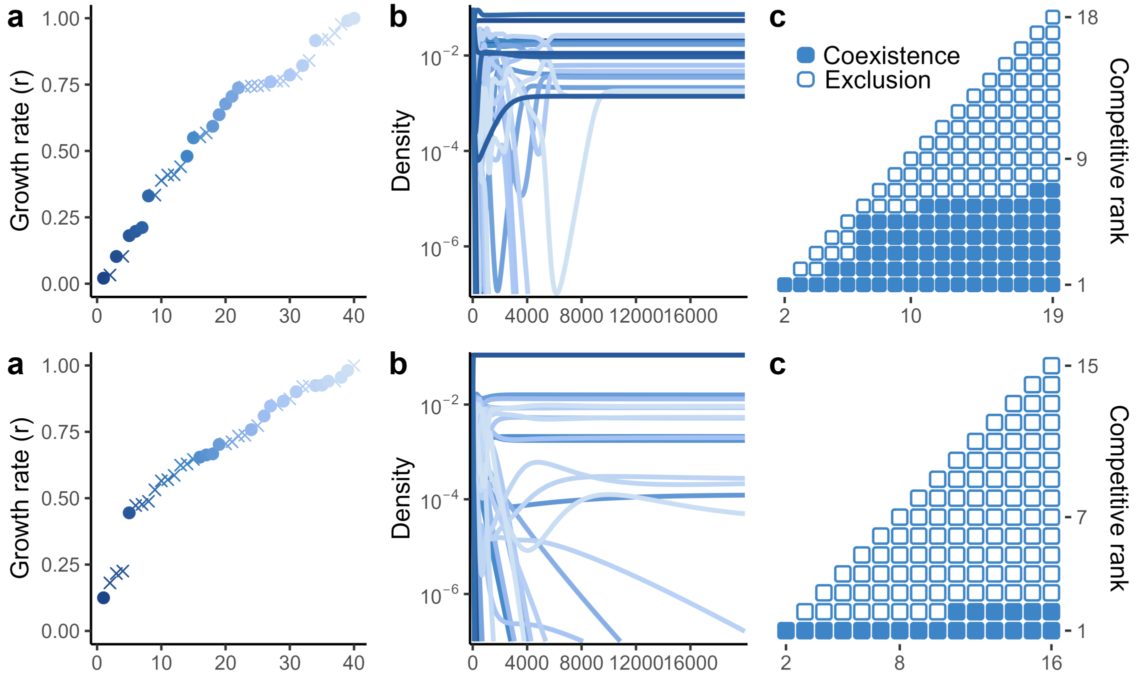


Figure S3: **Emergent coexistence in large, self-assembled communities in the competition-colonization trade-off model**. As in Fig. 3, main text, but model dynamics follow Eq. S32. **(a)** Growth rates are sampled independently from a uniform distribution on [0,1] and sorted so that $r_1 < r_2 < \dots r_n$. We translated growth rates into colonization and local extinction rates using $m = 1$ and $c_i = r_i + m$. **(b)** For an initial pool of 40 species, we numerically integrated the dynamics until a stable community of 19 species formed (21 species decline to extinction). In (a), persistent species are shown with filled circles, extinct species by crosses. **(c)** A majority of species pairs from the assembled community fail to coexist in isolation. **(d-f)** As in (a-c), except that growth rates are sampled from a triangular distribution (i.e., $P(r < \rho) = \rho^2$ on [0,1]), yielding a concave trade-off. Consequently, a smaller fraction of pairs coexist from the assembled community. In both cases the set of persistent species is very similar, but not identical to the growth competition trade-off model (Fig. 3, main text). The number of persistent species and coexistence fraction in the assembled communities are also very similar between the two models.

### C Emergent coexistence in random GLV communities

We found that even without an imposed trade-off, random (coexisting) GLV communities contained many non-coexisting species pairs and less intransitivity than expected by chance. The absence of known mechanisms for emergent coexistence (i.e., trade-offs or intransitive loops) led us to investigate factors that might drive pairwise exclusion alongside full community coexistence.

Based on our analysis of the trade-off models (above) and other studies of emergent coexistence (Saavedra *et al.*, 2017; Aguadé-Gorgorió & Kéfi, 2025), we hypothesized that widespread pairwise exclusion is explained by strong interspecific interactions and reconciled with full community coexistence by the presence of positive indirect (net) effects that arise when many species compete together. Our sampling procedure drew inter- and intraspecific interaction coefficients from the same (exponential) distribution, making it likely to find strong competitive interactions. We can further examine the effect of pairwise interaction strengths on pairwise coexistence by plotting, for each parameter set, the ratio of mean interspecific to mean intraspecific interaction strength against the coexistence fraction (among pairs). These results are shown in Fig. S4. Indeed, higher ratios of inter- to intraspecific competition are strongly associated with lower pairwise coexistence fractions. This result is consistent with well-known coexistence criteria for species pairs (Barabás *et al.*, 2016; Saavedra *et al.*, 2017).

Given that some of our simulated communities have mean interspecific interaction strengths comparable to or even exceeding mean intraspecific interactions, it is perhaps surprising that these communities coexist at all. Our sampling procedure was designed to find coexisting communities by selecting growth rates compatible with a positive (feasible) equilibrium and rejecting unstable parameter combinations. We can ask how the resulting ensemble of parameters, selected to allow coexistence, differs from the underlying distribution of parameters (without requiring feasibility or stability) as one way to understand the factors that allow the full communities to coexist. Specifically, we examined the proportion of indirect (net) effects that were positive, as a factor likely to promote coexistence in these communities. Indirect or net effects can be found by taking the inverse of the competition matrix $A^{-1}$. This matrix expresses the net effect of a press perturbation to one species on the equilibrium abundance of all others, as can be seen from the factor that equilibrium abundances are given by $x^\star = A^{-1}r$ (from Eq. 1). The signs of the elements $A^{-1}_{ij}$ tell us how the equilibrium abundance of a species $i$, $x_i^\star$, would change if the growth rate of another species $j$ was increased. Equivalently, the matrix $A^{-1}$ can be seen as summarizing the net effect of all interaction chains by which one species affects another (because $A^{-1}$ can be expanded as a Neumann series; see (Aguadé-Gorgorió & Kéfi, 2025)).

For each parameter set, we computed $A^{-1}$ and counted the fraction of interspecific (off-diagonal) effects that were positive. In Fig. S5, we compare this to the distribution of positive effects if we instead sample $a_{ij}$ i.i.d. from an exponential distribution and do not impose feasibility or stability. It is clear that conditioning on coexistence of the full community induces a shift toward a higher fraction of positive indirect effects.

In Fig. S6, we plot the transitivity metrics shown in the main text for other community sizes ($n = 4, 5, 7, 9$).

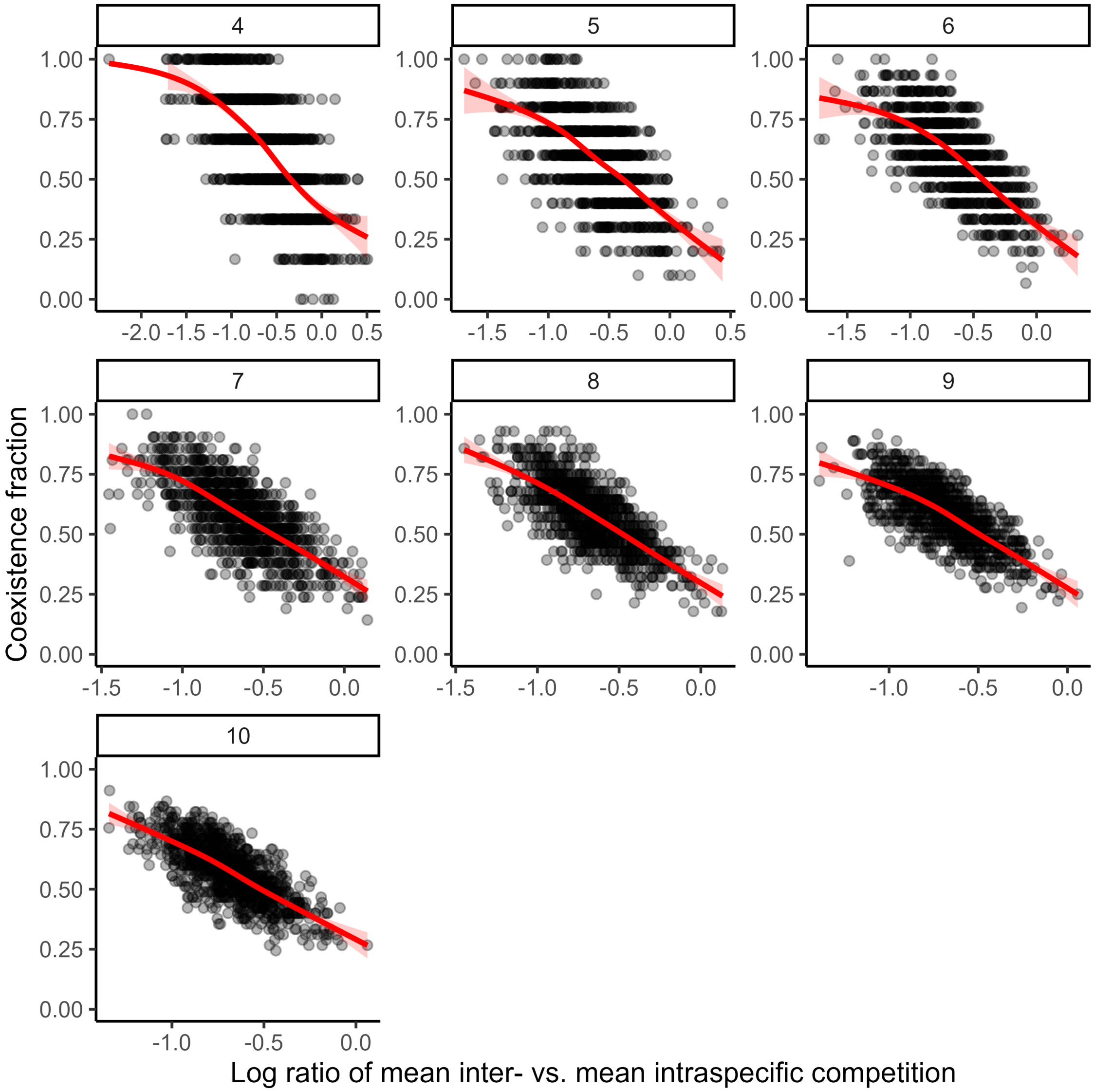


Figure S4: **Stronger interspecific competition reduces pairwise coexistence**. Each point represents one model community, with parameters obtained by random sampling conditioned on coexistence, as described in the text. The log ratio of mean inter- vs. mean intraspecific competition is computed as $\log\left[\left(\sum_{i,j\neq i} a_{ij}\right)/(n-1)\sum_i a_{ii}\right)\right]$. Smoothed conditional means (using geom_smooth from the ggplot2 package in R) are shown in red. Panels show different community sizes (number of species).

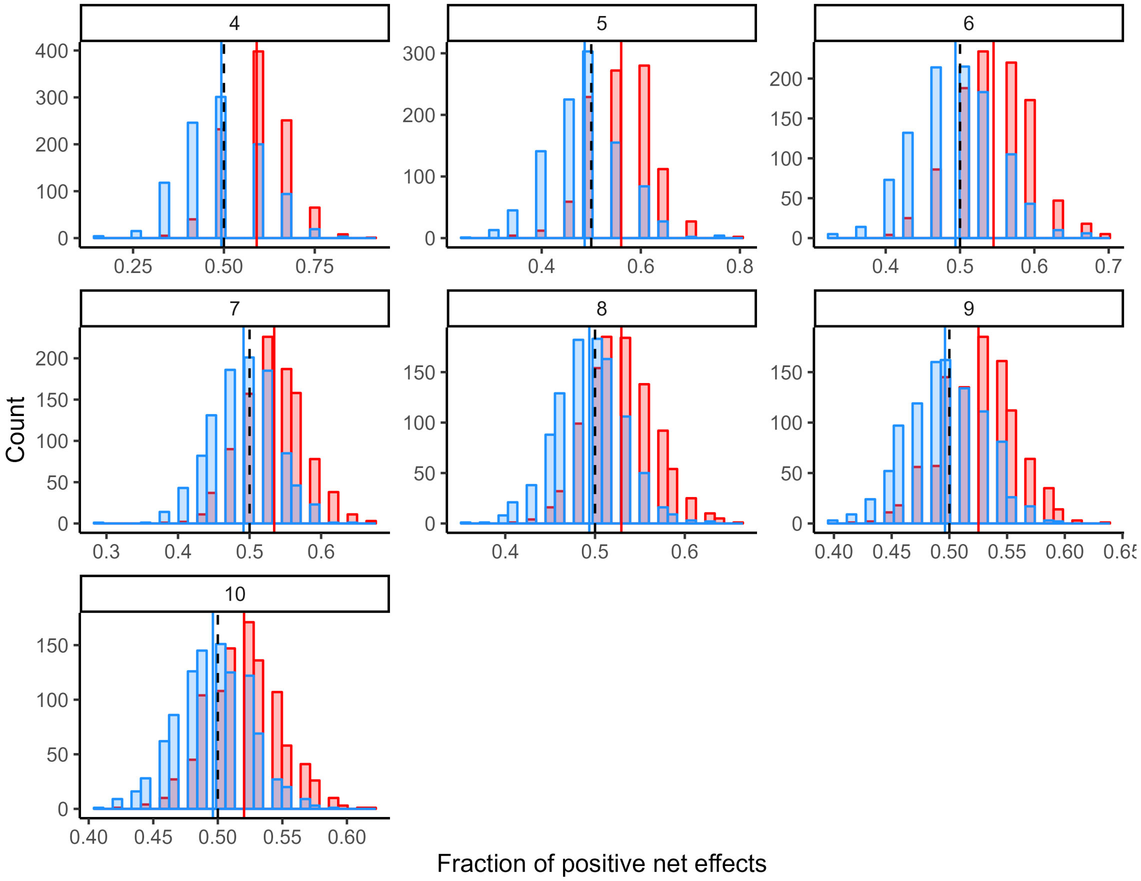


Figure S5: **Coexisting GLV communities have an excess of positive indirect effects**. The distribution of the fraction of positive indirect effects (positive off-diagonal elements of $A^{-1}$) across all model communities (conditioned on coexistence) is shown in red. A null distribution, obtained by inverting random matrices not conditioned on coexistence, is shown in blue. The mean for each distribution is indicated by a vertical line. The dashed line indicates equal proportion of positive and negative indirect (net) effects. Panels show different community sizes (number of species).

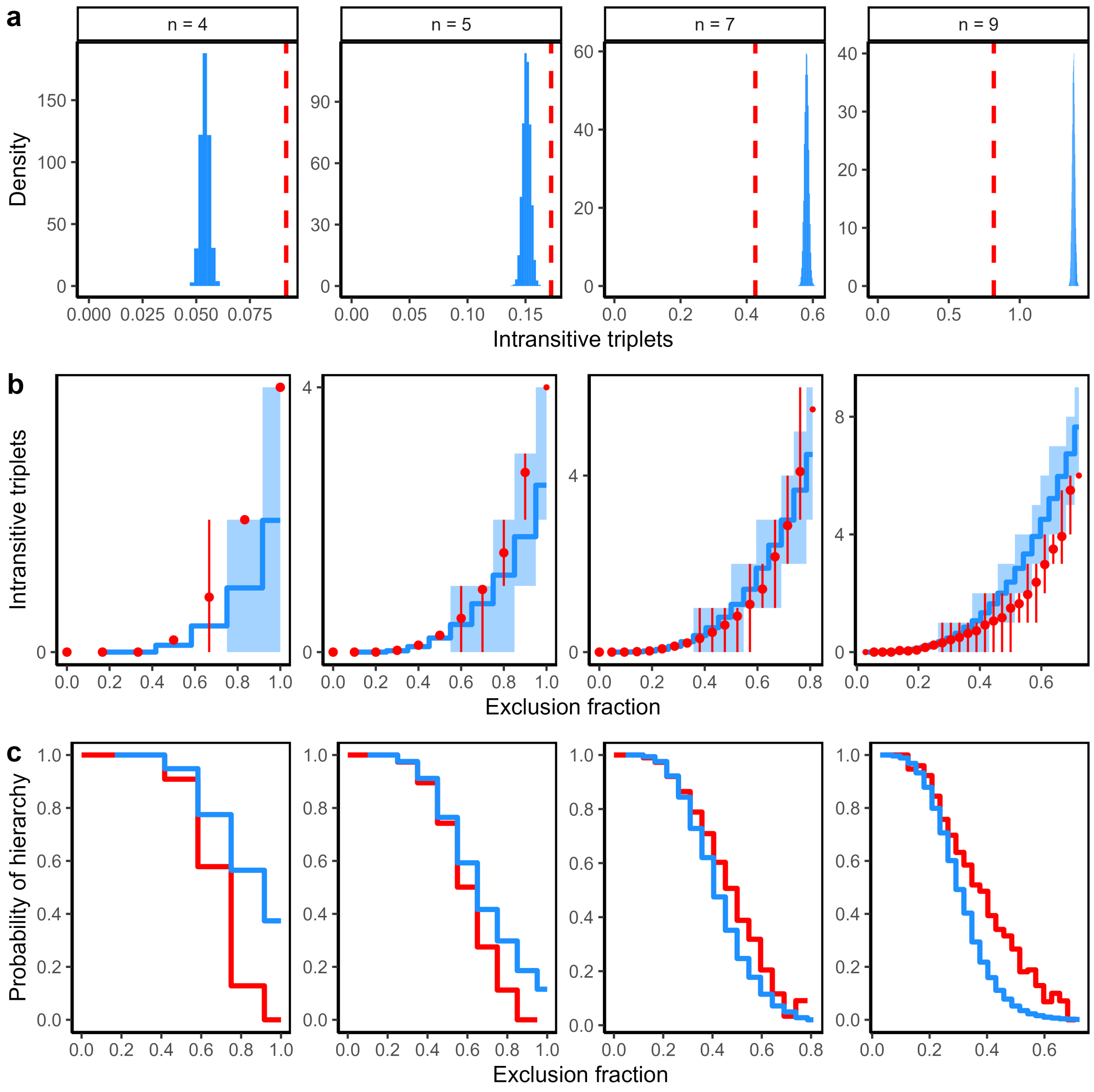


Figure S6: **Transitivity metrics in coexisting GLV communities,** $n = 4, 5, 7, 9$. As in Fig. 5, main text.
